# Mobilization of antibiotic resistance genes differ by resistance mechanism

**DOI:** 10.1101/2021.01.10.426126

**Authors:** Tue Kjærgaard Nielsen, Patrick Denis Browne, Lars Hestbjerg Hansen

## Abstract

The degree to which antibiotic resistance genes (ARGs) are mobilized by insertion sequence (IS) elements, plasmids, and integrons has a strong association with their likelihood to function as resistance determinants. This stems from genetic decontextualization where strong promoters often present in IS elements and integrons and the copy number effect of plasmids contribute to high expression of accessory genes. Here, we screen all complete bacterial RefSeq genomes for ARGs. The genetic contexts of detected ARGs are investigated for IS elements, integrons, plasmids, and phylogenetic dispersion. The ARG-MOB scale is proposed which indicates how mobilized detected ARGs are in bacterial genomes. Antibiotic efflux genes are rarely mobilized and it is concluded that these are often housekeeping genes that are not decontextualized to confer resistance through overexpression. Even 80% of β-lactamases have never, or very rarely, been mobilized in the 15,790 studied genomes. However, some ARGs are indeed mobilized and co-occur with IS elements, plasmids, and integrons. These results have consequences for the design and interpretation of studies screening for resistance determinants, as mobilized ARGs pose a more concrete risk to human health, especially under heterologous expression, than groups of ARGs that have only been shown to confer resistance in cloning experiments.

## Main

### Resistance determinants and context

Health and environmental challenges imposed by antibiotic resistance has sparked enormous research efforts into characterising genetic resistance determinants. Combined with broad availability of 2nd and 3rd generation sequencing technologies, studying the presence and prevalence of putative ARGs in the environment has become popular in recent years.

Antibiotic resistance remains a significant global issue despite numerous studies into understanding spread of genes via mobile genetic elements (MGEs) and devising and implementing mitigation strategies. But, most classes of putative ARGs are intrinsic to bacterial genomes and can be considered housekeeping genes. These do not confer resistance until they become ’decontextualized’ by mobilization^1–3^. Many putative ARGs have only been shown to provide resistance when cloned into cloning vectors with significant overexpression or with constitutive expression in mutants. Examples of mobilization events include (i) transfer from chromosomes to plasmids via transposons (Tn), (ii) integration with integrative conjugative elements (ICE), and (iii) capture by integrons as a gene cassette^4^. This aspect is often ignored in culture-independent studies using (meta)genomics and/or (q)PCR-based detection where resistance is rarely experimentally verified. Thus, screening the environment for ARGs may give impressions that “resistance is everywhere” or that widespread resistance predates the use of antibiotics^5–7^, although the native roles of found ARGs may not be related to antibiotic resistance^1, 2^.

### Native roles of ARGs

When coupling (meta)genomic predictions with culturing-based detection of resistant strains, it is often found that the two approaches do not agree^8–12^, partially attributable to the fact that gene expression is rarely considered in these studies^13^. Screening a genome for resistance markers against an ARG database often results in copious false-positive resistance predictions^10^. This issue is most pronounced for efflux-related markers where the specificity of prediction has been reported to be only 0.12. The balanced accuracy of resistance marker prediction against two widely used databases was only 0.52 and 0.66, showing that finding ARGs in a genome does not necessarily equate phenotypic resistance^14^. Almost all ARGs likely have native roles unrelated to resistance to clinical concentrations of antibiotics^1^. Many antibiotics are natural secondary metabolites, occurring at clinically insignificant concentrations, that are involved in inter-cellular communication, regulation of metabolism, and other housekeeping functions^15^. Putative ARGs have been found in and cloned from susceptible bacteria^8^, where they are simply performing their original non-resistance roles. For instance, efflux pumps^2, 16^, β-lactamases^17, 18^ and lipid A modifying proteins (MCR)^19^ are encoded by genes which have housekeeping functions but may be decontextualized to confer resistance.

Functional metagenomic approaches, essentially cloning fragmented DNA into expression vectors followed by screening for antibiotic resistance^6, 7, 20, 21^, has led to the identification of many putative ARGs. Such genes are obviously decontextualized in the experiment and their native roles may not be related to resistance. This has resulted in a problematic dissemination of resistance-related annotations in gene databases. Thus, sequence homology is a poor proxy for resistance and culture-independent techniques will often yield misleading results if genetic contexts of ARGs are not considered. With recent advances in long-read sequencing, high-quality metagenome-assembled genomes can be achieved^22^ and it is now possible to include consideration of the genetic context of ARGs.

### Classifying ARGs and their risk

The association between ARGs and MGEs have profound effects on phenotypic resistance^3, 4, 23–25^ and it has been argued, e.g. in the “RESCon” framework^26^, that multiple aspects should be included in risk assessment of ARGs^27, 28^. Here, we initiate the route to more accurate ARG predictions by categorizing associations between ARGs and MGEs in all completed RefSeq genomes. Decontextualization of ARGs is explored here by association with I) IS elements, II) plasmids, III) integrons, and IV) their dispersal across distinct genera.

## Results

### Screening RefSeq genomes for ARGs and data processing

The curated CARD database for ARGs was used to find putative ARGs in all completed bacterial genomes from the RefSeq database (n=15,790). 12,170 bp up- and downstream of predicted ARGs were analysed for IS elements and integrons, while the type of replicon (plasmid or chromosome) was also considered. For more details, see Methods and Supplementary Information (Supplementary Note 1). All databases are assumed to be heavily biased, especially towards human-associated bacteria of which many almost identical genomes have been uploaded to RefSeq, leading to overrepresentation of these compared to e.g. environmental bacteria. In order to ameliorate these biases (Supplementary Note 3), highly similar genetic loci with predicted ARGs (n=176,888) were clustered to 53,895 Clustered Resistance Loci (CRLs), representing 1,176 Antibiotic Resistance Ontology (ARO) terms from CARD (Fig. 1). This reduced the taxonomic Euclidean distance between CARD and RefSeq databases from 30.89 to 10.26, showing that many ARG loci in RefSeq are highly similar (Supplementary Note 4). Loci with efflux-associated ARGs were more compressed by clustering than all other types, indicating that these are more conserved and undergo fewer genetic rearrangements. The antibiotic efflux (AE) mechanism is the most abundant category and its CRL count is more than two times more numerous than the second-largest category, antibiotic inactivation (AI), although AI has over 3 times as many AROs as AE (Fig. 1).

**Fig. 1:**
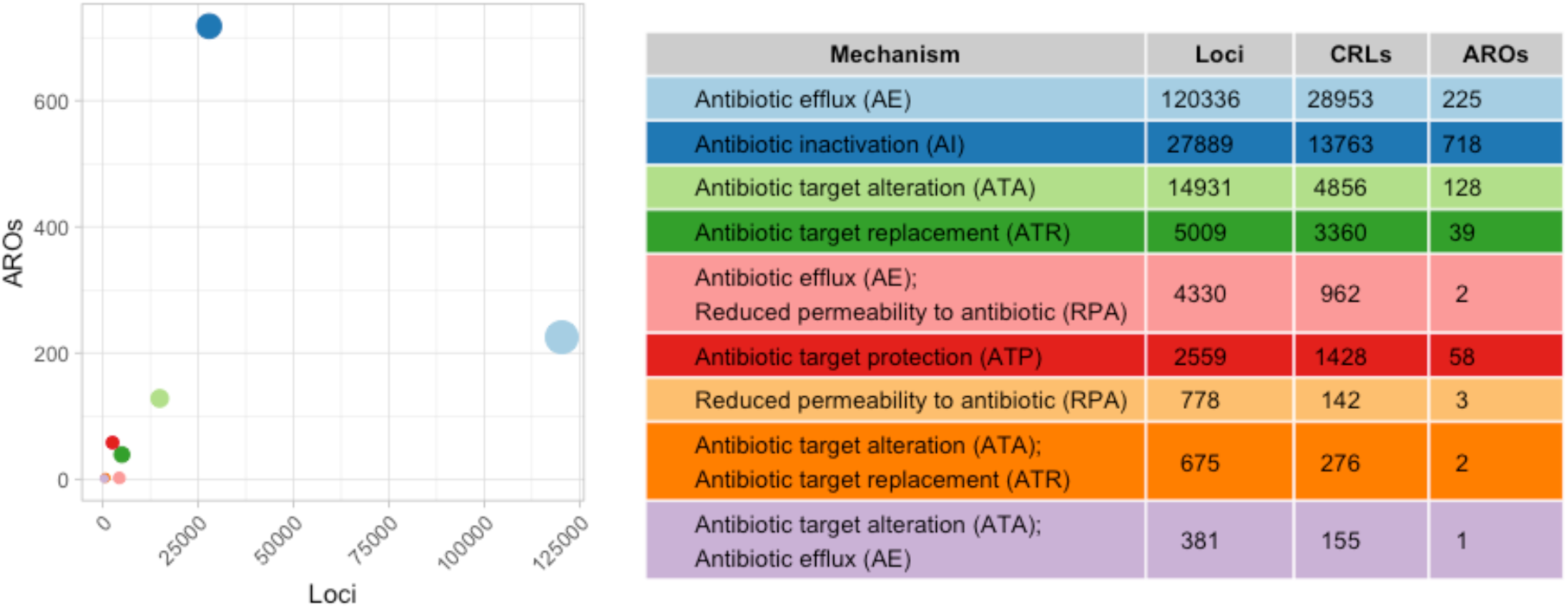
Overview of counts of ARG loci, Resistance Clustered Loci (CRLs), and CARD Antibiotic Resistance Ontology terms (AROs). In the plot, points are sized according to their CRL count (see table). Row colours in table correspond to their point colours in the left plot. The three hybrid mechanisms AE;RPA, ATA;ATR, and ATA;AE, as well as the low-ARO RPA mechanism, are excluded from some analyses, as they are not considered “main mechanisms”.

As expected, loci in human-associated genera were especially compressed by clustering, showing that these are indeed overrepresented in the RefSeq database (Supplementary Fig. 8).

Four mobilization parameter ratios were explored for each CRL (Fig. 2): A) replicon type, B) IS element-association, C) integron-association, and D) dispersal of CRL across genera (Simpson diversity). All parameters were calculated on a scale of 0-1, with 1 indicating that a CRL is highly associated with the given parameter. The mean of the four ratios is termed the ARG-MOB score and indicates how much genes of a given ARO are mobilized on a scale of 0-1. This is described further in later sections and Methods (Supplementary Fig. 9). Prophages in genomes are not explored for ARGs, since these are not thought to be common vectors^29^. Neither are ICEs explored, although they are likely important in resistance development^30^, since they are underexplored and may be difficult to predict in genomes.

**Fig. 2:**
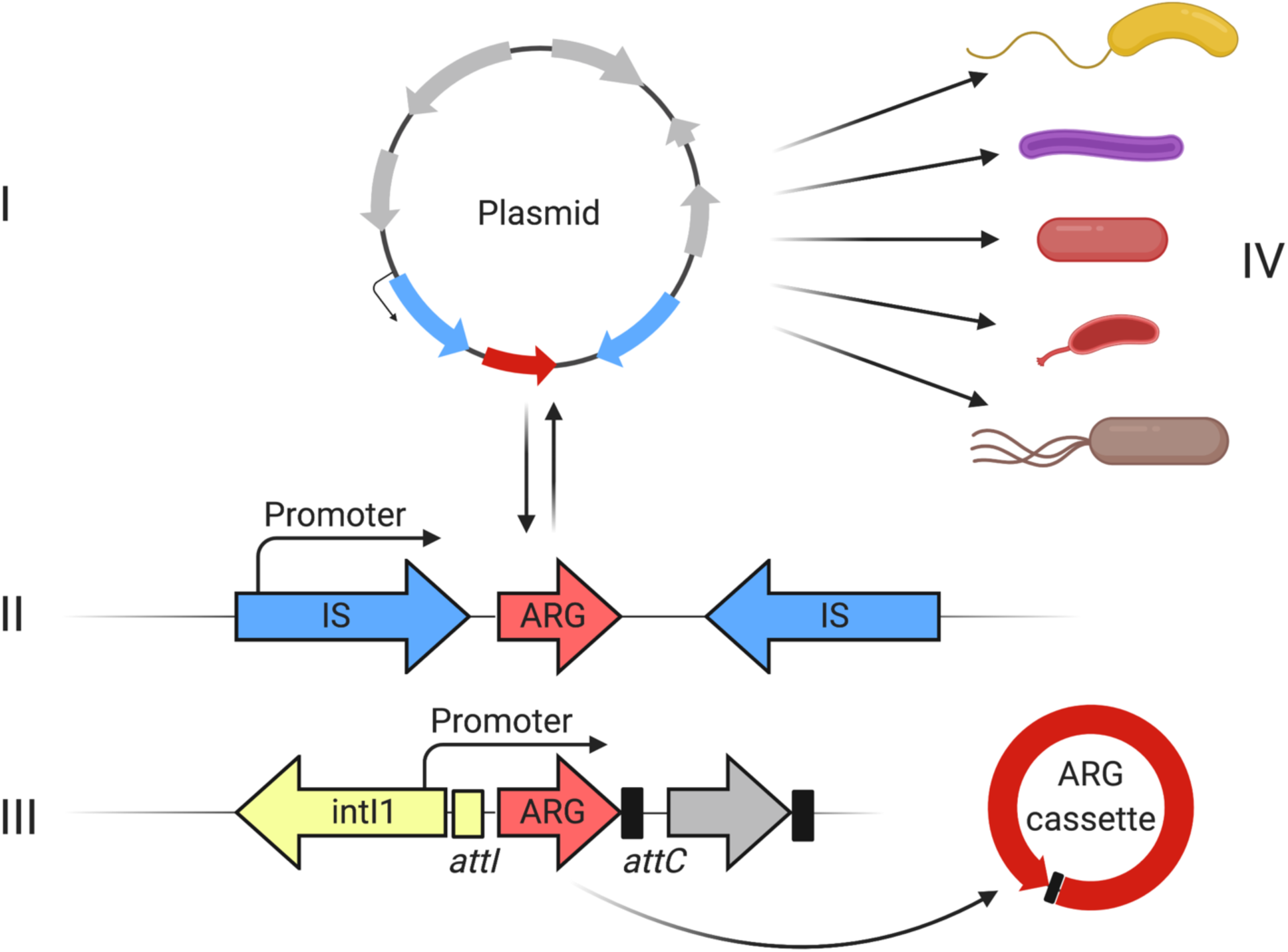
The four investigated mobilization parameters. I) classification of the replicon type that an ARG loci was found on, II) presence of one or more IS elements within 12,170 bp either up- or downstream of ARG, III) association of found ARG with integrons, and IV) the phylogenetic spread across genera, calculated by the Simpson diversity index. ARGs residing on plasmids can be rapidly spread horizontally and, in the case of multicopy plasmids, may be under heterologous expression. Many IS elements have either an internal promoter that can overexpress accessory genes or they may contain an outward-facing -35 component that can form a hybrid promoter, if the IS element is inserted close to a -10 box. If inserted as a gene cassette in an integron, the ARG is likely overexpressed by the common integron promoter. Furthermore, a gene cassette containing an ARG may form circular DNA molecules from the integron cassette array that can be shuffled to other locations. The final factor considered in this study with regards to mobilization of ARGs is the already observed phylogenetic dispersal of said ARGs across the genera represented in the RefSeq complete genomes database.

### Association of ARGs with IS elements and plasmids

Major resistance mechanisms (non-hybrid) are associated with IS elements and plasmids in varying degrees (Fig. 3). Furthermore, individual mechanisms were associated with different families of IS elements (Supplementary Note 5). AE AROs generally have very low IS and Replicon ratios, which indicates that ARGs of this mechanism are rarely mobilized by either IS elements or plasmids. Only a handful of AE AROs have both high IS and high Replicon ratios, including ARO3002693 (transposon-encoded *cmlA1* chloramphenicol exporter), ARO3003836 (*qacH* subunit of fluoroquinolone exporter), and ARO3000165 (tetracycline efflux pump *tetA*). Many AE AROs contain a large number of unique CRLs, as is also reflected by AE CRL count in Fig. 1. Therefore, AE ARGs are rarely associated with either IS elements or plasmids. Distances, in terms of nucleotides, between ARGs and IS elements are larger for AE AROs than for other mechanisms, indicating that efflux ARGs are more “loosely” associated with IS elements than other mechanisms (Mann-Whitney U-test (MWU) P < 0.01; Supplementary Note 6). Contrary to AE, the AI mechanism has many AROs that have been mobilized by both IS elements and plasmids, but also AROs that are hardly mobilized at all (Fig. 3). With some exceptions, antibiotic target alteration (ATA) AROs have low IS and Replicon ratios while also exhibiting a low number of unique CRLs, indicating that ATA CRLs are conserved and often not decontextualized. On the other hand, antibiotic target replacement (ATR) AROs are more mobilized by IS elements and plasmids (Fig. 3).

**Fig. 3:**
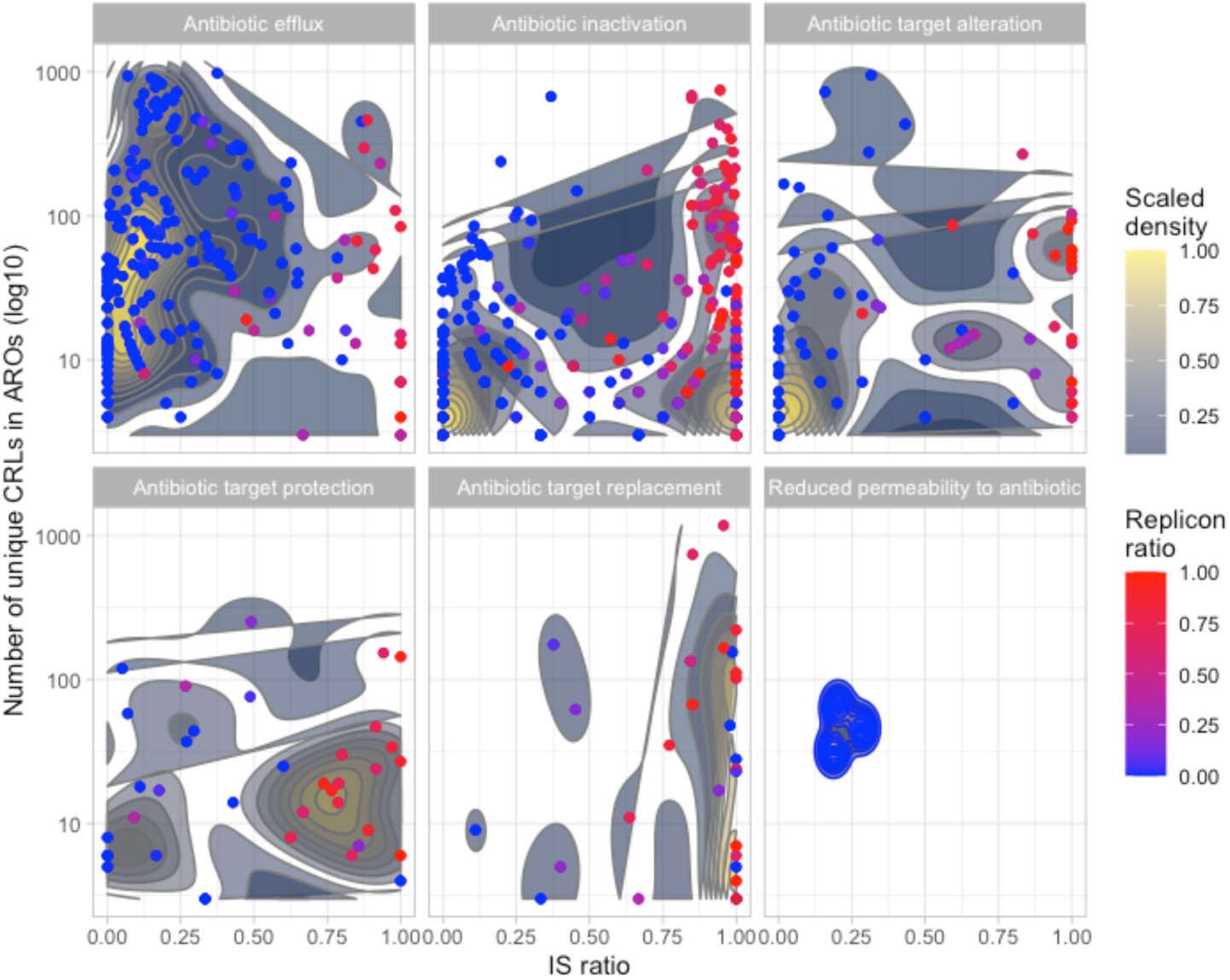
Density plots of IS ratio against the number of unique clustered CRLs in a given ARO category, represented by individual points. Plots are divided into the individual mechanisms and are colored according to the Replicon ratio, where a high ratio (red) indicates that an ARO is more often found on plasmids and a low ratio (blue) indicates that an ARO is more on chromosomes. Density estimates are calculated with two-dimensional kernel density estimation, as implemented in the stat_density_2d function under the ggplot R package. The hybrid mechanisms are not included.

Putative ARGs are more decontextualized in clinically relevant genera (Supplementary Figs. 12-14). As expected from database biases (Supplementary Figs. 6-8), Proteobacteria harbour 88.18% of unclustered ARG loci (Supplementary Fig. 2) and Proteobacteria have a higher median IS ratio than Firmicutes, Actinobacteria, and Bacteriodetes (Supplementary Note 7; MWU test; p < 0.05). Within Proteobacteria, multiple families with clinically relevant members, such as *Enterobacteriaceae*, are highly associated with ARGs mobilized by IS elements and plasmids.

Phyla whose members are more associated with the environment, such as Actinobacteria and Bacteriodetes, have lower median IS ratios than Proteobacteria. Generally, diving into specific families and genera shows that ARGs in human-associated bacteria are more mobilized than in others (Supplementary Table 3) and persistent fixation of mobilized ARGs are likely a consequence of human interference with pathogenic bacteria^23^. This is described in more detail in Supplementary Note 7.

### Integron-association varies across ARG classes

Under selective pressure for resistance, ARGs may be decontextualized into integrons, where a strong promoter confers overexpression of said ARGs, resulting in phenotypic resistance. Using IntegronFinder^31^ on CRL sequences, 3,723 ARGs were identified as gene cassettes in integrons or clusters of *attC* sites lacking integron-integrases (CALIN). The most abundant major mechanism was AI with 2,684 unique CRL occurrences. ATR and AE were found in association with integrons in 694 and 310 CRLs, respectively (Supplementary Fig. 15). Interestingly, canonical sulfonamide resistance genes associated with Tn*402* Class 1 integrons^32^, *sul1-4,* were not the most frequent submechanism associated with integrons and was here found associated with integrons in only 123 unique CRLs out of 2,017 total *sul1-4* CRLs. Trimethoprim resistance *dfr* genes associated with Class 2 integrons and Tn*7* transposon^32^ were here found in high abundance in association with integrons. The most abundant submechanism was the antibiotic inactivation ANT(3’’) category for nucleotidylylation of aminoglycosides at the hydroxyl group at position 3’’ with 1,009 CRLs associated with integrons. ANT genes are often found in association with integrons^33^ and subgroups of ANT genes are discussed further in Supplementary Note 8. OXA-9 and OXA-1 β-lactamases are found in integrons in 98.51% and 63.90% of the 67 and 277 CRLs, respectively, emphasising that these ARGs are of concern (Supplementary Table 4).

From these observations, it is evident that screening an environment with PCR for e.g. sulfonamide resistance genes will result in prediction of many unmobilized genes that are not overexpressed through an integron promoter. On the other hand, OXA β-lactamases are often associated with integrons and thus likely to confer resistance, but nevertheless these results emphasize the importance of considering the genetic contexts of ARGs.

### Mobilization assessment based on four parameters

Inspired by previous work^1^, we calculated a mobilization scale for each ARO, termed the ARG-MOB scale, which is based on IS and plasmid ratios, integron-associations, and dispersal across genera, as calculated using Simpson’s diversity index. The ARG-MOB ratio is calculated as the mean of the four mobilization parameters (MOB) and ranges from 0 to 1 with 1 representing very high mobilization, signified by very high IS and plasmid ratios, frequent association with integrons, and a wide phylogenetic dispersal across genera. Fig. 4 shows the MOB parameters and ARG-MOB scale. For each parameter, boxplots with MWU test results are also shown.

**Fig. 4:**
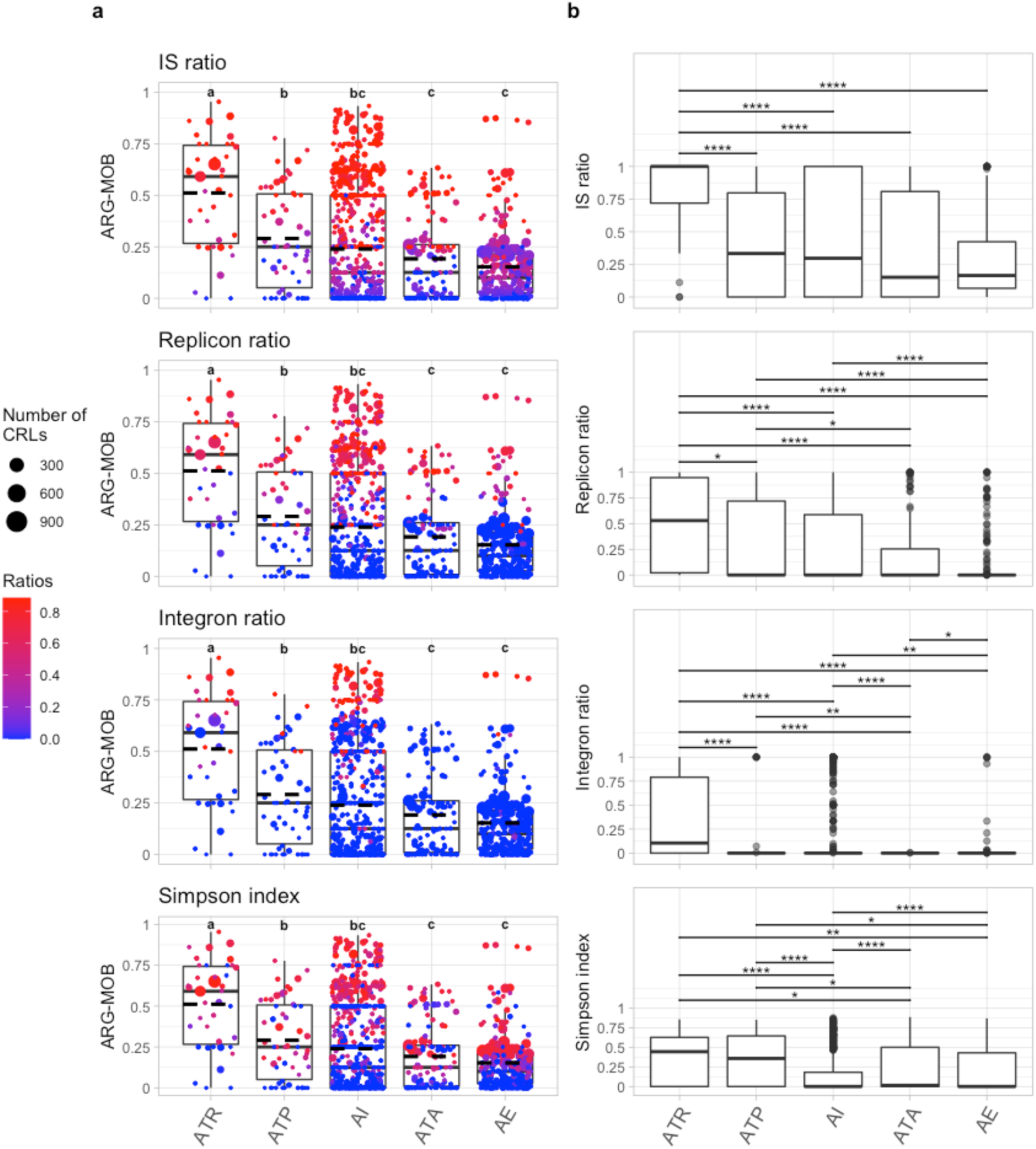
ARG-MOB scores of major resistance mechanisms defined from the four MOB parameters. **a**, ARG-MOB score of AROs by major mechanism. Each point indicates a specific ARO and the size of the point corresponds to the number of unique CRLs in that ARO. Each of the four plots shows one of the individual MOB parameters as coloured gradients of the points. Points are horizontally jittered but placed identically between the four plots in the left column. Mean is shown with dashed lines. Boxes indicate first and third quartiles (25% and 75% of data) and horizontal line in box shows the median. Whiskers extend to 1.5 * of the interquartile ranges. Letters above boxplots indicate significant differences between mechanism populations (Mann-Whitney U-test with FDR correction; P < 0.05). **b**, Boxplots of each mobilization factor per major mechanism. Outliers are shown as grey dots. Above boxplots, bars indicate significant differences in distribution between mechanisms (Mann-Whitney U-test with FDR correction). Only significant differences are displayed (*: p < 0.05;**: p < 0.01; ***: p <0.001; ****: p <= 0.0001).

The median ARG-MOB per major mechanism is highest for ATR (p < 0.0001), while AE has a low median ARG-MOB but not significantly different from ATA and AI groups. Antibiotic target protection (ATP) and AI groups are not significantly different (Fig. 4a).

### Efflux genes are rarely mobilized

The AE mechanism has the lowest median ARG-MOB (although only significantly lower than ATR and ATP), which is reflected by median Replicon, IS, and Integron ratios that are lower than most other groups (Fig. 4), i.e. efflux genes are rarely mobilized by these MGEs. It is therefore likely that most identified AE ARGs are housekeeping efflux pumps located in conserved loci of chromosomes, with a few exceptions (e.g. multidrug efflux genes *oqxAB*^34–38)^. This warrants caution to include considerations of genetic context when screening environments for efflux-associated ARGs.

The highest median ARG-MOB mechanism, ATR, is characterized by AROs with a high degree of mobilization by IS elements and plasmids (Fig. 4). The high ARG-MOB ATR AROs are furthermore strongly associated with integrons and are taxonomically more wide-spread than ATP, AI, and AE groups. Likewise, some ATP AROs are highly mobilized and widespread but they are to a lesser degree associated with integrons. Generally, ATR is significantly more associated with integrons than other categories, although the median of AI is higher than ATA and AE.

AROs of AI mechanism are the least phylogenetically dispersed but are instead conserved within few genera, as indicated by the lowest median Simpson index. Possibly, many genes and/or proteins under the AI mechanism only function in specific genera, whereas those of other mechanisms can function in wider ranges of genera. While there certainly are AI AROs that have been mobilized by plasmids, transposons, and integrons, there are many others that have not been decontextualized (Fig. 4). All major mechanisms have exceptions in the form of AROs with elevated ARG-MOB, as evaluated on all four parameters, although ATA, ATP, and AE have few or no AROs with ARG-MOB higher than 0.75.

### The ARG-MOB scale proficiently describes decontextualization of putative ARGs

The four MOB parameters all correlate significantly with each other, showing that they covary and are appropriate for calculating the ARG-MOB scale (Fig. 5a,b). Hierarchical clustering of AROs from all five major mechanisms in a heatmap shows apparent mechanism-specific profiles of ARG-MOB scores, as well as each of the four MOB parameters (Fig. 5c).Two major branches are formed from clustering: I) a high-ARG-MOB branch dominated by AI, as well as other individual AROs from other mechanisms and II) a low-MOB branch mostly populated by AE AROs.

**Fig. 5:**
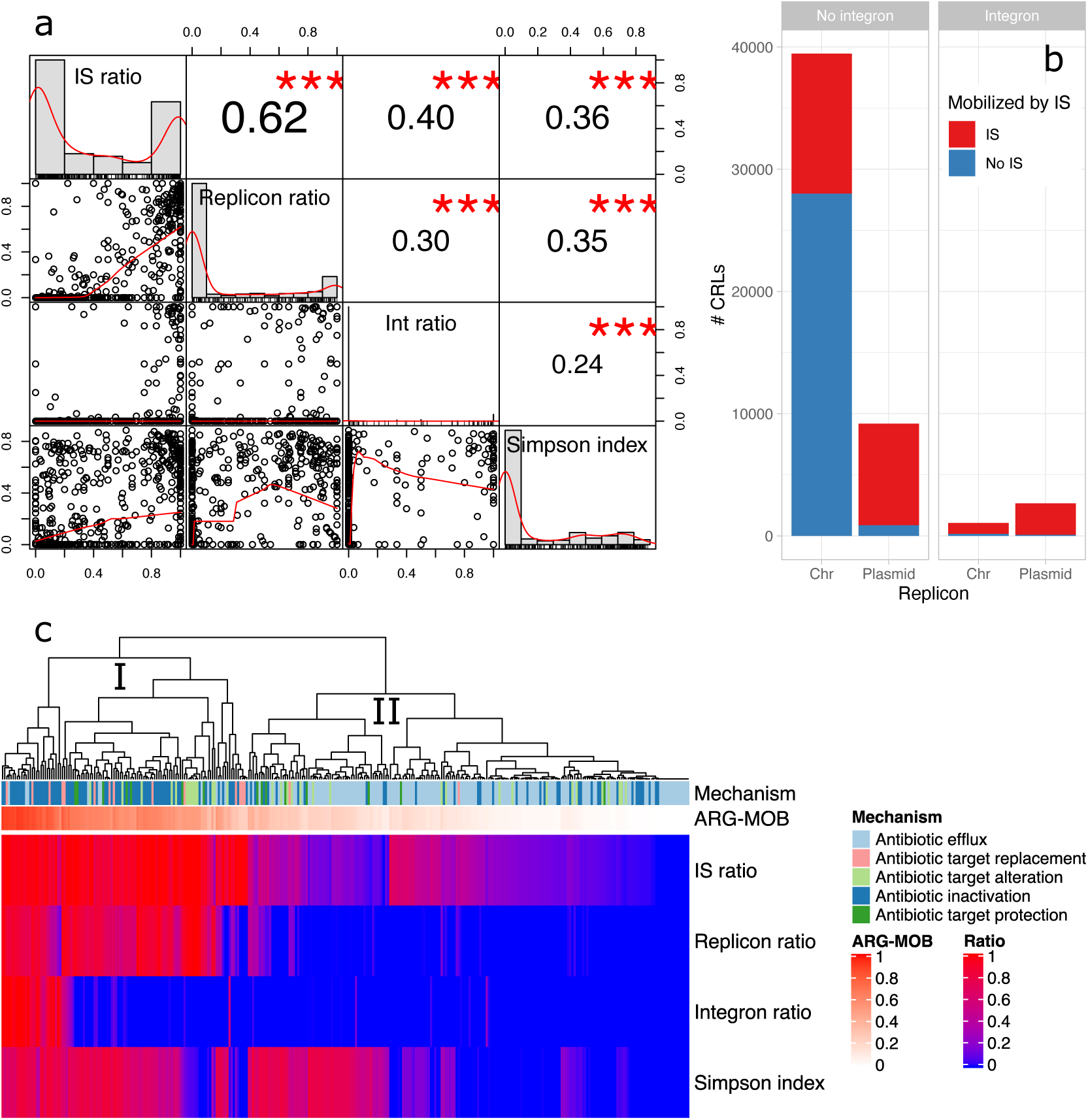
Correlation and co-occurrence of MOB parameters. **a**, Pearson correlation coefficients between MOB parameters. Scatter plots between pairwise MOB parameters are shown in lower left corner. The diagonal shows histograms of distribution of each MOB parameter. The values in the upper right corner show the Pearson correlation coefficients with significance levels (***: P < 0.001). **b**, Barplot of mobilization of unique CRLs by IS elements, plasmids, and integrons. **c,** Heatmap of highly abundant AROs with at least 20 CRLs. The dendrogram shows clustering of the AROs, based on the four MOB parameters, and was calculated using standard parameters in the ‘ComplexHeatmap’ package (complete hierarchical clustering on Euclidean distances).

The highest correlation coefficient is seen for IS-Replicon ratios, showing that ARGs placed on plasmids are likely mobilized by IS elements prior to insertion on plasmids (Fig. 5a,b). The second highest correlation is found between IS and integron ratios, indicating that ARGs, found as gene cassettes in integrons, are likely to have been mobilized (as part of integrons) by IS elements (Fig. 5a,b) which has been often reported and discussed^4, 24, 32^. To a lesser degree, ARGs found on plasmids are correlated with integrons.

Perhaps not surprisingly, the Simpson diversity index correlates positively with IS, replicon, and integron ratios (Fig. 5a). This shows directly that highly mobilized ARGs and those found in integrons are also likely to be phylogenetically widespread. On the basis of these correlations, we conclude that the ARG-MOB ratio proficiently describes decontextualization of putative ARGs. Pearson correlation coefficients and MGE co-occurrences were also calculated per mechanism (Supplementary Note 9).

### Some AROs are highly divergent in mobilization

Many AROs can be defined as either highly mobilized or only to a very little degree. Still, some AROs have a very large spread from their ARG-MOB score, showing that they are most often sitting unmobilized on a chromosome, but have one or times been mobilized and widely dispersed (Supplementary Figs. 22,23). This is exemplified by the efflux pump encoded by genes *oqxAB* (ARO3003922-3)^34, 35^ (Supplementary Note 10). These genes are found on essentially all *Klebsiella pneumoniae* chromosomes where they do not confer resistance unless highly overexpressed^36–38^. However, they were shown to be mobilized by IS elements on plasmid pOLA52 where they confer resistance to multiple antibiotics^34^. The *oqx* AROs show high spread across their mean IS and Replicon ratios (*oqxA* has ratios of 0.35 and 0.16, respectively), showing that mean ratios are low due to *Klebsiella* chromosomes but that there are many outliers due to variants in *Escherichia* and *Salmonella* that are only found mobilized by IS elements and usually on plasmids (Supplementary Fig. 22; Supplementary Table 5).

Outliers from the mean of IS and Replicon ratios can also be considered per genus instead of ARO, in order to highlight that ARGs in some genera are much more mobilized than in others. For example, efflux pump genes in *Shigella* are much more associated with IS elements compared to the global average, but they are not found on plasmids more than on average (Supplementary Fig. 23). Likewise, many AI ARGs are found more on plasmids in *Escherichia*, *Salmonella*, *Klebsiella*, *Citrobacter*, and *Enterobacter* than their respective average placements per ARO. Other genera including *Proteus*, *Pseudomonas*, *Acinetobacter*, and *Morganella* tend to have some AI ARGs more located on chromosomes than the given ARO average, indicating that chromosomes in these genera may be considered reservoirs of genes with potential as resistance determinants. This highlights the complexity of the ARG issue and emphasises the importance of considering genetic context of ARGs before predicting resistance.

### Defining ARG-MOB categories

Smoothed kernel density estimates of AROs and their ARG-MOB values are shown in Fig. 6a per mechanism and cumulatively for all mechanisms. The following five ARG-MOB groupings were defined computationally: *Very low* (ARG-MOB = 0), *Low* (0 < ARG-MOB < 0.182), *Medium* (0.182 < ARG-MOB < 0.378), *High* (0.378 < ARG-MOB < 0.681), *Very high* (ARG-MOB > 0.681). These definitions are largely the same when estimating per mechanism individually (Supplementary Fig. 24). Numerically, AI has the highest number of *High* and *Very high* ARG-MOB AROs (144 and 54, respectively), while ATR has the highest percentage of *High* and *Very high* ARG-MOB AROs with these categories representing 64% of ATR AROs (Fig. 6b).

**Fig. 6:**
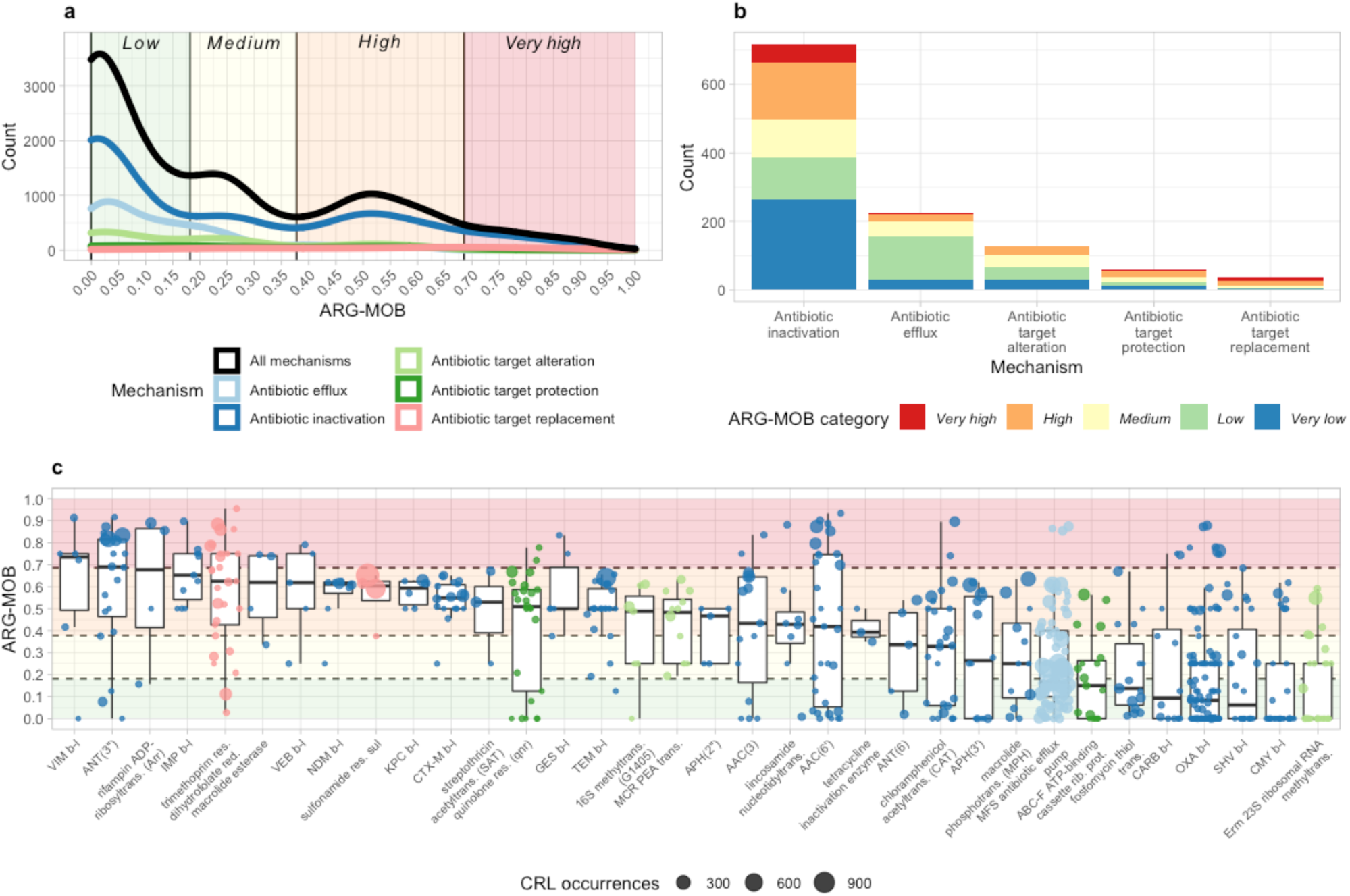
Density of ARG-MOB categories and distribution per major resistance mechanism. **a**, Count density of ARG-MOB per mechanism. Only major non-hybrid mechanisms are shown. ARG-MOB categories were defined computationally by finding valleys in “All mechanisms” density estimates, except for the border between High and Very high where no valley is present. These calculations are described in more detail in Methods and Supplementary Fig. 9. **b**, Count of each ARG-MOB category per mechanism. **c**, All submechanisms that have AROs with *High* or *Very high* ARG-MOB scores. Points are coloured by major resistance mechanism as in Fig. 6a and sized according to the number of CRLs of the given ARO. The background is coloured similarly to fig. 6a, representing *Low*, *Medium*, *High*, and *Very high* categories. The *Very low* category is for ARG-MOB = 0 and does thus not have a colour in the graph.

### *High* ARG-MOB AROs correspond with high-risk ARGs

*High* and *Very high* ARG-MOB AROs (Fig. 6c) are mainly ARGs that were initially identified in very serious and resistant pathogens where they indeed confer resistance. Vice versa, many low ARG-MOB AROs have only been shown to confer resistance when placed on high-expression cloning vectors but not in any natural wild-type isolate. A few examples are described below and in Supplementary Note 11. A comprehensive table for all 1,176 AROs can be found as an interactive table in Supplementary Table 6.

For all major mechanisms, many AROs are classified as *Very low* or *Low* ARG-MOB (Fig. 6c) and ATA does not have any *Very high* ARG-MOB AROs, while ATP has two (ARO3002803 and 3002801; quinolone resistance genes *qnrVC6* and *qnrVC4*). AI has many AROs with *High* and *Very high* ARG-MOB, which include infamous β-lactamases, aminoglycoside nucleotidyltransferases (ANT), and others (Fig. 6c). With a median ARG-MOB of 0.73, the Verone integron-encoded metallo-β- lactamase (VIM) is the β-lactamase with the highest median ARG-MOB. There are three VIM β-lactamase AROs, of which ARO3002271 has the highest ARG-MOB of any AI ARO at 0.91. In RefSeq complete genomes, it is only found inserted in integrons, and is located close to IS elements and on plasmids in 95% of the CRLs found (n=21). It is dispersed across 6 unique genera for a Simpson index of 0.75 (*Pseudomonas*, *Salmonella*, *Escherichia*, *Klebsiella*, *Citrobacter*, and *Enterobacter*). VIM-1 was isolated from a multiresistant *E. coli* from a patient. It was inserted in a Class 1 integron and found on a conjugative plasmid^39^. It has since been seen in multiple *Enterobacteriaceae*, typically in association with MGEs, and is globally spread^40^.

The highest ARG-MOB ATR AROs belong to the trimethoprim resistant dihydrofolate reductase *dfr* submechanism. The ARO3003013 within this submechanism has the highest ARG-MOB of any ARO at 0.95. A Class 1 integron with *dfrA15* is widespread in *Vibrio cholera* isolates in Africa and was found on a conjugative plasmid^41^. It is the ARO with the highest ARG-MOB, since it was only found to be associated with IS elements, integrons, and plasmids (all ratios = 1). It has a Simpson index of 0.82 and the 7 CRLs are dispersed across 6 genera (*Vibrio*, *Salmonella*, *Enterobacter*, *Leclercia*, *Klebsiella* and *Escherichia*).

Based on examples of high and low ARG-MOB AROs (Supplementary Note 11), a pattern emerges that high ARG-MOB AROs were originally isolated from and identified in already virulent, pathogenic bacteria that had indeed been tested as resistant, often by acquisition of new genetic material. On the other hand, low ARG-MOB AROs were generally identified in susceptible bacteria and/or only shown to cause resistance when cloned into vectors with strong gene expression. This warrants caution when choosing ARGs of interest in either targeted (q)PCR screening or metagenomic sequencing of environmental samples.

## Discussion

Ideally, the genetic context is always included in ARG predictions, since even the highest ARG-MOB scoring genes will have representatives that are not decontextualized and may not confer resistance. This study documents how genes from even the most mobilized categories of ARGs can be found unmobilized on chromosomes. Our results clearly demonstrate that the validity of using PCR-based screening to assess the abundance and distribution of putative ARGs is questionable at best (e.g. for antibiotic inactivation ARGs) and almost certainly miscalling others (antibiotic efflux ARGs). This is in agreement with previous studies finding discrepancies between genotypic and phenotypic resistance predictions^8–12^, with especially efflux-related markers producing a high number of false-positive predictions^14^. Therefore, it is necessary to consider the genetic context of putative ARGs when trying to assess the relevance of ARGs^1, 3, 26^. This could be achieved by applying PCR primers that target regions spanning both an ARG and an associated MGE^42^. For more accurate ARG calling, metagenomic sequencing using long-read platforms is a prerequisite to enable the detection of ARGs and their genetic contexts.

## Methods

### Databases and ARG prediction

All code for data processing was written in BASH scripts and statistics and plotting were primarily done in RStudio. All scripts and a searchable table of results (Supplementary Table 6) are available at https://github.com/tueknielsen/ARG-MOB. The details of the methods are described below.

All complete bacterial genomes (15,790 entries with 16,785 chromosomes and 14,280 plasmids) were downloaded from RefSeq on Dec. 12 2019 using the ncbi-genome-download tool v0.2.11 (available from https://github.com/kblin/ncbi-genome-download). In order to ensure uniform prediction of genes across all bacterial genomes, Prodigal^43^ (v2.6.3) was used to predict genes from nucleotide sequences and write corresponding amino acid sequences from all RefSeq genomes. Since Prodigal first trains itself based on the input sequence, gene prediction was performed on subsets of each genus present in RefSeq genomes. Per genus, two rounds of Prodigal were performed with the -meta flag enabled in the second run, as it predicts some genes that are missed in single genome mode and vice versa. Results from the “single” and “meta” gene predictions were combined and redundant annotations found with both methods were merged.

Several ARG databases and tools for predicting ARGs have been produced, including CARD^44^, ARDB^45^, MEGARes^46^, ResFinder^47^, SARG^48^, ARG-ANNOT^49^, DeepARG-DB^50^, ARG-miner^51^, FARME^52^, and others. Some are discontinued while others receive updates occasionally. The CARD database is large, actively updated, well-curated, and widely used. Furthermore, it makes use of ontology terms (Antibiotic Resistance Ontology: ARO) that allow for the grouping of resistance genes according to resistance mechanisms. Because of these advantages over other databases and the essential role of ontology terms, the CARD database was used in this study. The “protein homolog” models from CARD were used here, since they do not contain resistance determinants that are based on mutations. The main resistance mechanisms defined in the CARD database are “antibiotic efflux” (AE), “antibiotic inactivation” (AI), “antibiotic target alteration” (ATA), “antibiotic target protection” (ATP), “antibiotic target replacement” (ATR), and the less abundant “reduced permeability to antibiotic” (RPA). A few additional categories exist that are hybrids of two of the above mechanisms, but there are very few entries of these in CARD and are for most of the analyses not considered.

The Comprehensive Antibiotic Resistance Database (CARD v3.0.7) was downloaded and only the protein homolog model was used in this study, excluding resistance determinants related to sequence variants (e.g. SNPs). DIAMOND^53^ blastp was used to identify putative ARGs in all RefSeq genomes. For blastp against the CARD database, both query and subject coverages were set to a minimum of 80%, while E-value cutoffs were set to 1e-10, to limit the rate of spurious hits. For each query protein from all RefSeq genomes, only the single best CARD match was kept.

The CARD auxiliary tool, RGI^54^, for predicting ARGs in (meta)genomes uses curated blastp bitscore cutoffs unique to every ARG protein in the CARD database. The same bitscore cutoffs were applied here, with the exception that hits with bitscores lower than the RGI cutoff were included if they had an identity score and a query coverage of at least 80%. These hits were included in order to keep more putative ARG hits from environmental bacteria that are not clinically relevant, since it is assumed that CARD and other ARG databases are biased towards genes that reside in anthropogenically relevant strains. Blastp hits with bitscores above the RGI cutoff were also only kept if query coverage was at least 80%. The effects of these filters are further described in Supplementary Information (Supplementary Figs. 1,2,3; Supplementary Note 1).

Data tables were imported into R for statistics and visualization using the packages ggplot2^55^, dplyr^56^, tidyr^57^, gridExtra^58^, ggpubr^59^, ggExtra^60^, reshape2^61^, knitr^62^, kableExtra^63^, vegan^64^, PerformanceAnalytics^65^, ComplexHeatmap^66^, RColorBrewer^67^, DT^68^, rstatix^69^, tidyverse^70^, broom^71^, and plotly^72^. All statistical tests on rank-sums of groupings were performed with unpaired Mann-Whitney U-tests (MWU) with Benjamin-Hochberg FDR correction for multiple testing.

### Extracting the genetic context of ARGs

The average length of composite and unit transposons were calculated based on 449 entries in the Transposon Registry^73^. This average (12.17 kbp) was used as the maximum allowed distance between an ARG and an IS element for classifying an association (Supplementary Table 1). However, since ARGs in transposons can be on either strand relative to the transposase, IS elements are identified within 12.17 kbp of an ARG in both directions. This enables searching for transposons of up to 24.34 kbp (plus the length of the identified ARG), which would include 77.73% of the 449 composite and unit transposons in The Transposon Registry^73^ (Supplementary Fig. 4).

For all filtered blastp ARO hits, up to 12,170 bp both up- and downstream of the hit were extracted from the respective RefSeq replicon using the faidx command from Samtools^74^ (v1.9-166-g74718c2).

If an ARG was found within 12.17 kbp of either terminus of a replicon, only sequence until the terminus was extracted and not continued from the other end of the sequence, since entries in RefSeq complete genomes may not be actually complete, due to low sequencing coverage regions stemming from e.g. GC-bias in sequencing^75^. Loci were categorized according to the ARO of the identified ARG. There are 9 ARO major mechanism categories of which three are less abundant “hybrids” merged by two other categories. The 6 non-hybrid categories Antibiotic efflux (AE), Antibiotic inactivation (AI), Antibiotic target alteration (ATA), Antibiotic target replacement (ATR), Antibiotic target protection (ATP), and Reduced permeability to antibiotic (RPA) are here considered the main categories and are the ones mainly investigated in this study. The mechanism “Reduced permeability to antibiotic” is only represented by three ARO categories and is excluded from some statistical analyses.

### IS elements in ARG loci and 16S rRNA as control

IS elements in ARG loci were predicted using DIAMOND blastp against the ISfinder database^76^, as implemented in Prokka^77^ (v1.14.0). The same E-value cutoff for IS annotations, as Prokka applies during gene annotation (1e-30), was used here and the minimum query coverage accepted was 90%. Only the top IS hit for each query protein was kept, since multiple “good” hits to distinct IS families may occur per query. The distance between a given putative ARG and its closest IS neighbour within 12.17 kbp in either direction (if any) was calculated without considering the coding strand of the genes. ARGs not within 12.17 kbp of an IS element were not considered when calculating the mean ARG-IS element distances.

Since 16S rRNA genes are not expected to be often mobilized by IS elements, the distance between 16S rRNA genes and IS elements were explored in all complete RefSeq bacterial genomes, in order to assess how many “false-positive” ARG-IS associations are expected to be identified using the 12.17 kbp distance cutoff (Supplementary Note 2). 80,141 16S rRNA genes in 15,790 strains were predicted using barrnap^78^. Of these, 94.61% did not have identified IS elements within 12,170 bp in either direction, which can be seen as analogous to a 95% confidence interval for predicting association between ARGs and IS elements.

### Clustering ARG loci to remove redundancy

Extracted loci with putative ARGs were grouped based on the CARD ARO category of the loci ARGs. In order to remove redundancy from database bias (Supplementary Notes 3 and 4; Supplementary Figs. 6,7) towards oversampling of e.g. almost identical *E. coli* chromosomes, extracted loci were clustered with USEARCH^79^ (v11.0.667_i86linux64) (Supplementary Fig. 8 and Supplementary Table 2). Per ARO group, sequence loci were clustered into what we refer to here as Clustered Resistance Locus (CRL) using the “-cluster_fast” command with the criteria that sequences in a cluster are at least 99% similar over at least 90% of the length (both target- and query coverage) and only the single best hit was allowed per sequence. The “-sort length” flag was also enabled to sort loci by length before clustering, since loci vary in length (sum of 12.17 kbp up- and downstream plus an ARG of varying length). This ensures that loci of identical length (with the exact same ARG) are merged into the same CRLs. For each CRL, the centroid sequence was used as representative sequence for downstream analyses.

### Integron prediction

Integrons and cassette arrays were predicted using IntegronFinder^31^ using the centroid CRL sequences as input. IntegronFinder can predict complete integrons including gene cassettes, In0 elements where only integrase is present, and CALINs (Cluster of *attC* sites Lacking Integrase nearby). All three classes of integrons are included in the analyses and no distinction is made, since a putative ARG observed in e.g. a CALIN has been previously associated with an integron and may still be in related strains.

### MOB metrics and the ARG-MOB scale

Four main metrics (Fig. 2), or ratios (0 to 1), of mobilization were calculated per ARO that aim to quantify just how mobilized groups of ARGs are. These four ratios are A) the Replicon ratio B) the IS ratio C) the Integron ratio and D) the phylogenetic spread of an ARO across distinct genera, quantified by the Simpson diversity index. Pearson correlation coefficients between MOB metrics were calculated.

For each ARO category, the number of CRLs with and without identified IS elements were counted and the IS ratio was derived where an IS ratio of 1 indicates that all CRLs belonging to a given ARO have an IS element within 12,170 bp either up- or downstream of the ARG. Vice versa, an IS ratio of 0 indicates that none of the CRLs in an ARO have IS elements in proximity. Similarly, the Replicon ratio was calculated per ARO based on the CRLs’ location on either plasmids or chromosomes. A Replicon ratio of 1 means that all CRLs in a given ARO are of plasmid origin and a 0 means that all CRLs are from chromosomes. The Integron ratio indicates how many CRLs are inserted in integrons per ARO. For measuring the taxonomic distribution of each ARO category, the Simpson diversity index (range 0 to 1) was calculated per ARO using unclustered sequences and the genera they were identified in.

The ARG-MOB scale (0-1) represents the mean of the four MOB metrics described above and serves as a ranking scheme to evaluate how the degree to which members of an AROs have been mobilized. Based on the smoothed kernel density estimates of all ARG-MOB scores, groupings were made to categorize AROs by their ARG-MOB score. An ARG-MOB score of 0 indicates that ARGs of the given ARO were not once found to be mobilized in the RefSeq genomes and a score of 0 is thus categorized as *Very low*. Valleys in the density distribution of ARG-MOB scores were used to computationally pinpoint thresholds between ARG-MOB categories. The *Low* group ranges ARG-MOB score from 0.0 to 0.182, the *Medium* group ranges from 0.182 to 0.378, *High* ranges from 0.378 to 0.685, and *Very high* ranges from 0.685 to 1.0. For the *Low-Medium* and *Medium-High* cutoffs, the low point in valleys was used to define values but no apparent valley is present between *High* and *Very high*. Instead, a linear model was fitted to the right-side slope of the *High* peak and another fitted to the approximately linear data range starting at ARG-MOB score 0.7. The intersection between the two linear models (0.685) was used as the cutoff between the *High* and *Very high* groups (Supplementary Fig. 9).

## List of abbreviations

ARG: Antibiotic Resistance Gene
MGE: Mobile Genetic Element
Tn: Transposon
IS: Insertion Sequence
AE: Antibiotic Efflux
AI: Antibiotic Inactivation
ATA: Antibiotic Target Alteration
ATR: Antibiotic Target Replacement
ATP: Antibiotic Target Protection
RPA: Reduced Permeability to Antibiotic

## Supplementary information

Note: Supplementary information starts with information related to Methods and is followed by information related to Results. Supplementary references are listed separately from main article.

### Supplementary Note 1: Filtering DIAMOND blastp CARD hits

By default, all blastp hits with bitscores exceeding the per-ARG-curated RGI bitscore-cutoffs are accepted. A ratio (bitratio) is calculated by dividing bitscores with the RGI cutoffs where a bitratio of less than 1 indicates that the blastp hit has a lower bitscore than the RGI ARG cutoff. However, as can be seen in Supplementary Fig. 1, there are many blastp hits that have high percentage identity and high query coverage, although their bitscores are below 1 and would thus be discarded if only RGI cutoffs are considered. Considering the database biases described in the article and in further detail below, it is likely that these high-similarity but low-bitratio hits are actually true ARG homologs in strains that are not related to those highly abundant in the CARD database. To include these hits, another filter was introduced where hits with bitratio lower than 1 are still included if their % identity and % query coverage are above 80% (Supplementary Fig. 1).

**Supplementary Fig. 1.**
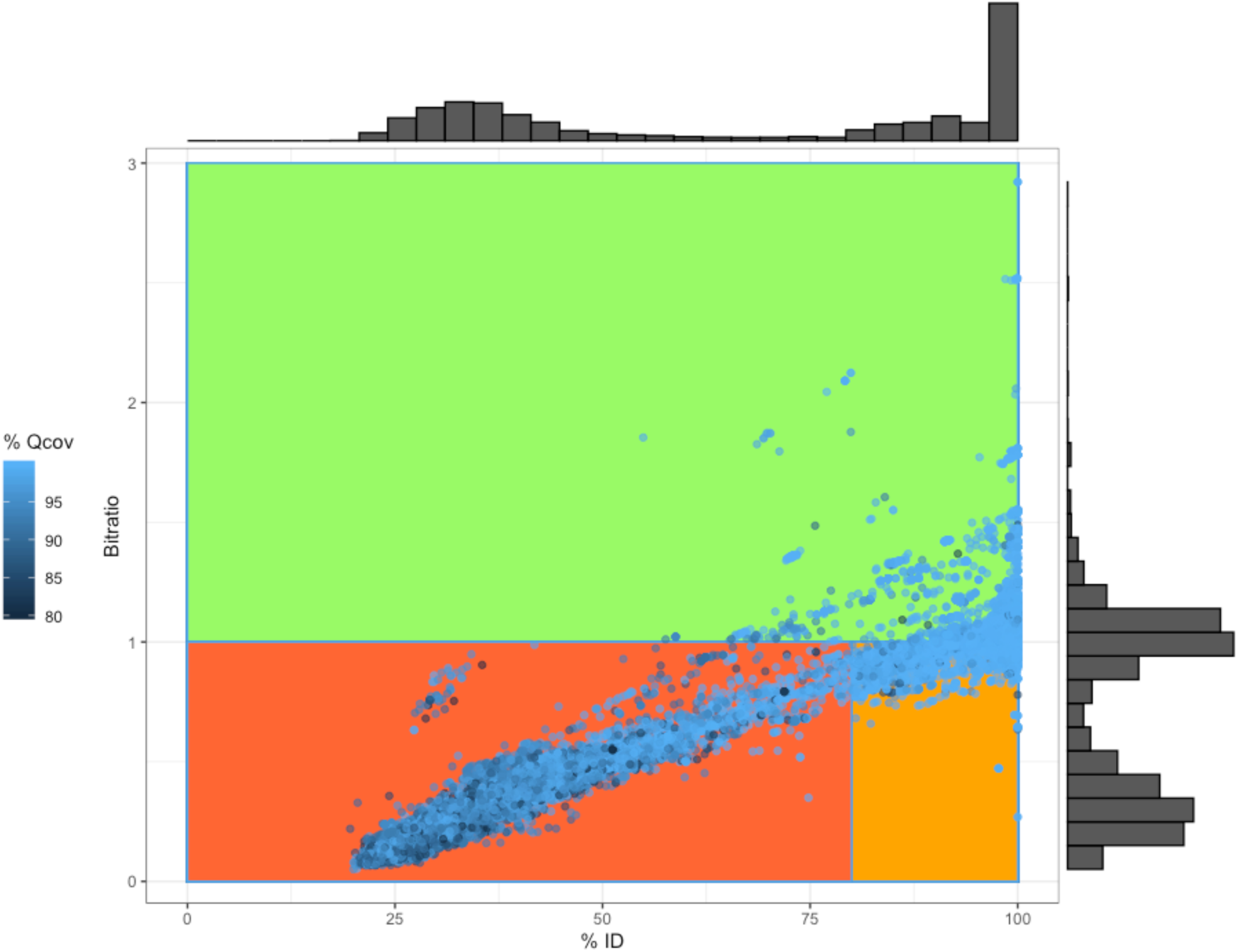
Distribution of DIAMOND blastp hits using all proteins from RefSeq complete genomes as queries against the CARD database. Before applying filters, the search cutoffs were at least 80% query coverage and an E-value of 10E-10. Subsequently, hits were filtered based on ARG- specific bitscores (curated by CARD/RGI) and by minimum 80% ID. Hits passing the RGI bitscore cutoff are locateds in the green area, while hits passing the custom 80% ID cutoff are located in the orange area. Hits not passing either filter are located in the red area. Bitratio on Y-axis is calculated by dividing individual bitscores by the curated bitscore-cutoffs from CARD/RGI. Histograms on the outside of the plot show the distribution of both %ID and bitratio.

Blastp hits passing the filters were investigated for phylogenetic distribution. By also including blastp hits that are more than 80% identical to a CARD protein, we include additional 61,620 hits on top of the 115,268 hits passing the RGI bitscore cutoffs. The major taxonomic orders are mostly equally included by the RGI bitscore filter and the 80% ID filter (Supplementary Fig. 2), with the biggest order being *Enterobacterales* that constitutes 25% and 49% of blastp hits passing the ID and RGI filter, respectively.

**Supplementary Fig. 2.**
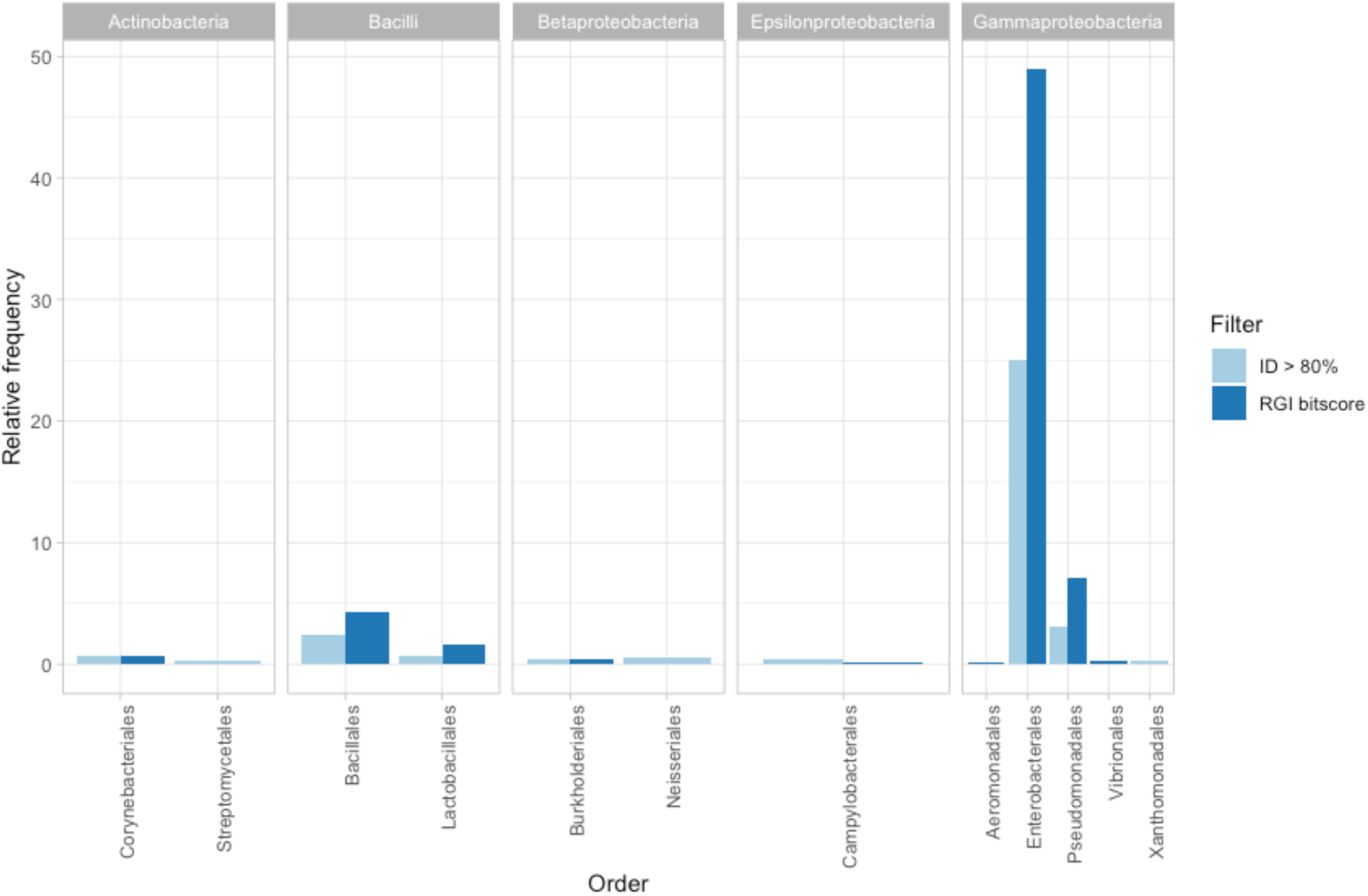
Major bacterial orders of ARG blastp hits passing either of the defined filters. On the Y-axis, the percentage of the total hits passing the respective filter is shown. Major orders are defined as orders that constitute more than 0.2% of the total data. In this plot, blastp hits can be in either ID filter or RGI filter, but not both (either bitscore > threshold or bitscore < threshold but ID > 80%).

However, the minor bacterial orders are not as equally distributed in RGI and ID filters as the major orders (Supplementary Fig. 3). Specifically, the orders *Micrococcales*, *Pseudonocardiales, Rhodospirillales*, *Aeromonadales*, *Alteromonadales*, *Legionellales*, and *Vibrionales* are passing the ID filter more than the RGI bitscore cutoff. This shows that including hits passing the additional ID filter expands the scope of this study to incorporate more environmental bacteria.

**Supplementary Fig. 3.**
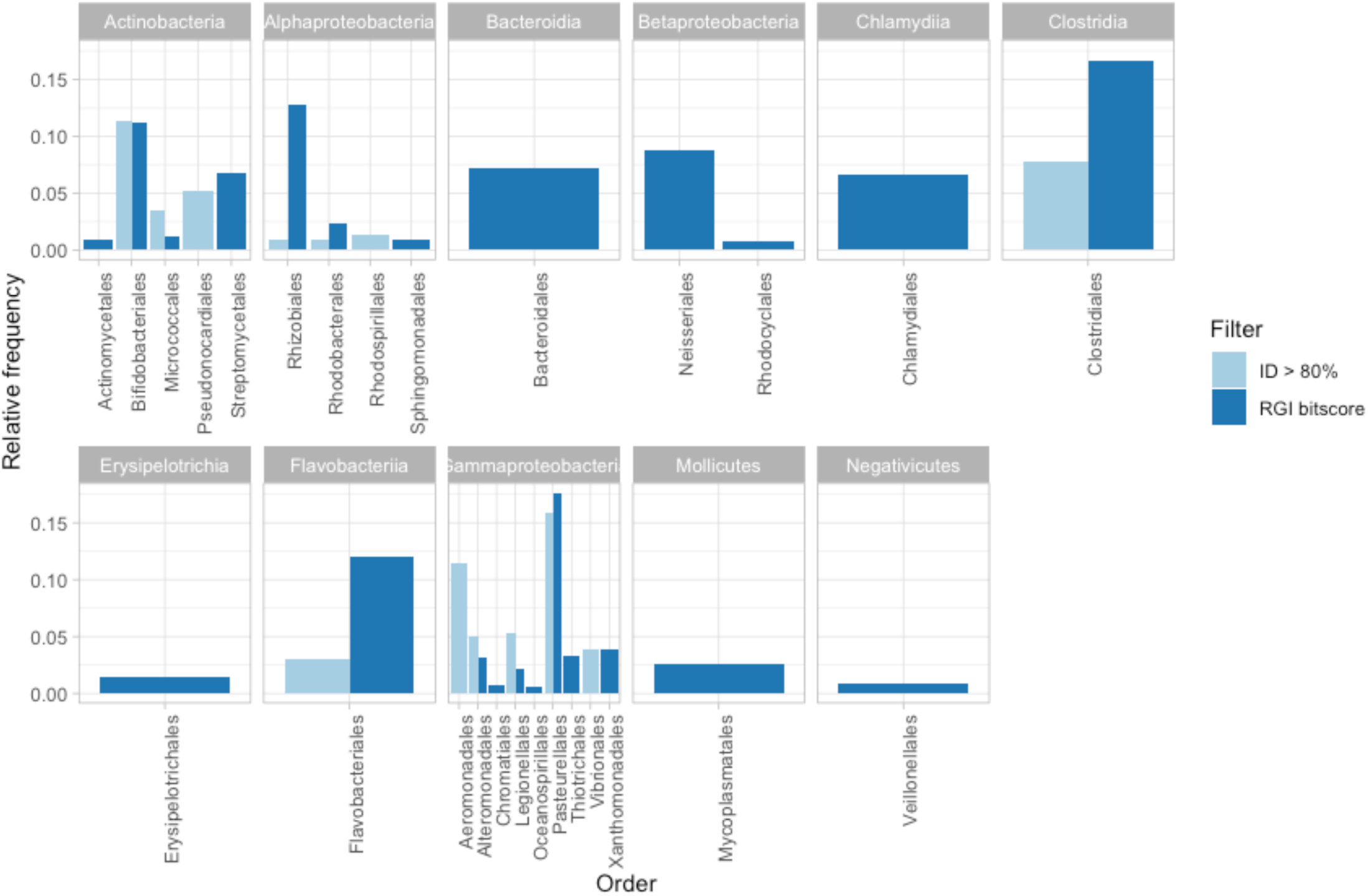
Minor bacterial orders of ARG blastp hits passing either of the defined filters. On the Y-axis, the percentage of the total hits passing the respective filter is shown. Minor orders are defined as orders that constitute less than 0.2% of the total data but only orders with more than 10 hits are shown here. In this plot, blastp hits can be in either ID filter or RGI filter, but not both (either bitscore > threshold or bitscore < threshold but ID > 80%).

#### Mean lengths of composite and unit transposons

**Supplementary Fig. 4.**
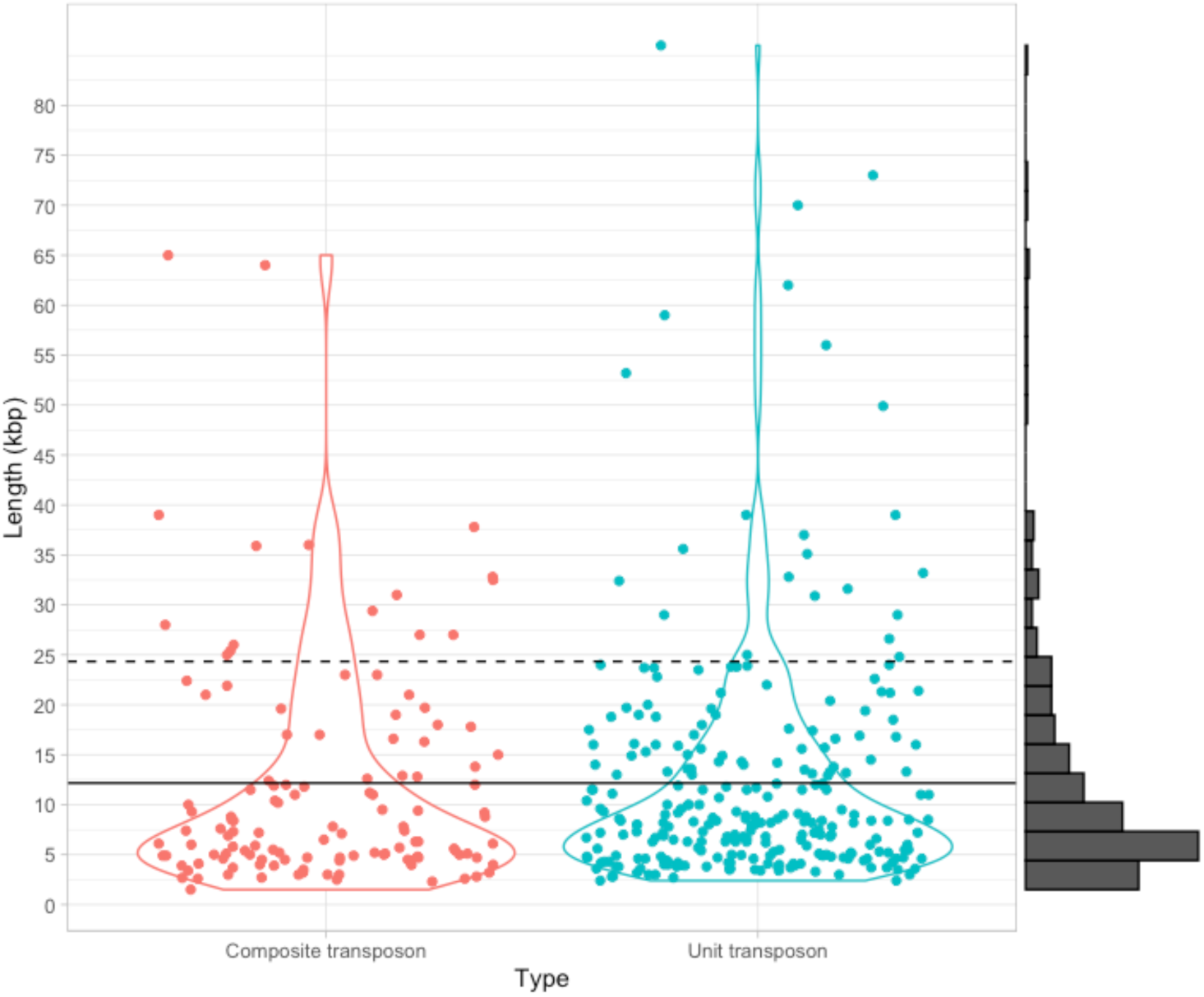
Violin plot of sequence lengths of 313 unit transposons and 136 composite transposons from The Transposon Registry1. The mean length (12.17 kbp) for both transposon types is shown as a solid line, and the maximum length of a genetic region with a putative ARG (24.34 kbp) is shown with a dashed line. A histogram of both length distributions combined is shown to the right. The individual mean lengths are 12.37 and 11.74 kbp for unit and composite transposons, respectively.

**Supplementary Table 1.**
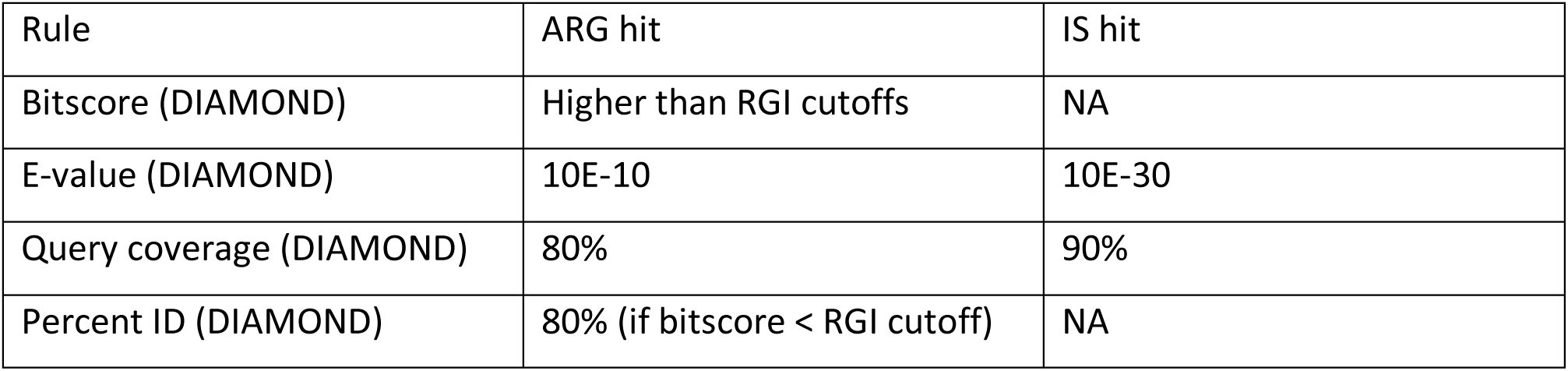
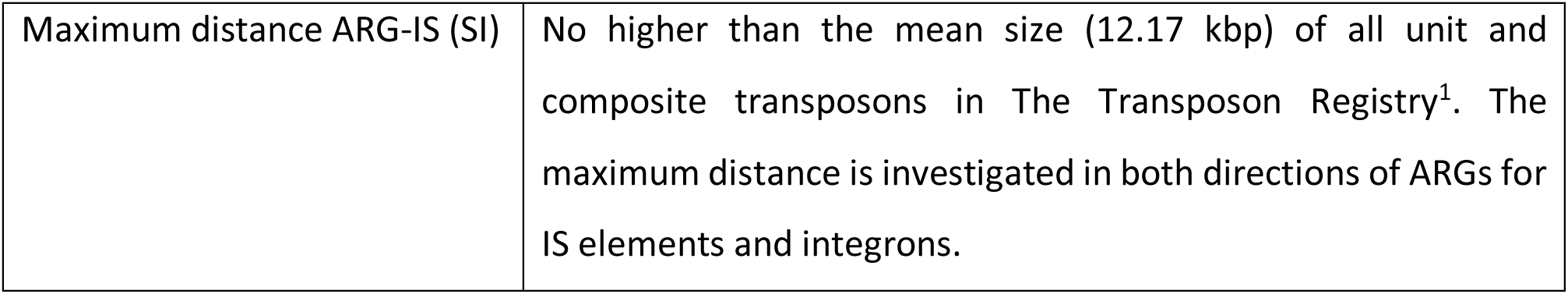
Cutoffs and search criteria for ARG and IS element prediction.

| Rule | ARG hit | IS hit |
| --- | --- | --- |
| Bitscore (DIAMOND) | Higher than RGI cutoffs | NA |
| E-value (DIAMOND) | 10E-10 | 10E-30 |
| Query coverage (DIAMOND) | 80% | 90% |
| Percent ID (DIAMOND) | 80% (if bitscore < RGI cutoff) | NA |

|  |  |
| --- | --- |
| Maximum distance ARG-IS (SI) | No higher than the mean size (12.17 kbp) of all unit and composite transposons in The Transposon Registry<sup>1</sup>. The maximum distance is investigated in both directions of ARGs for IS elements and integrons. |

### Supplementary Note 2: Distance between all 16S rRNA genes and closest IS elements

Using barrnap^2^, 80,141 16S rRNA genes were identified in 15,790 strains in RefSeq complete genomes (mean 5.08 16S genes/genome). Only IS elements within 100 kbp of 16S rRNA genes were considered. Out of the 80,141 16S rRNA genes 4,480 had one or more IS elements within 12.17 kbp, corresponding to 5.59% (Supplementary Fig. 5). This gives an acceptable accuracy of 94.61% when predicting associations between ARGs and IS elements.

**Supplementary Fig. 5.**
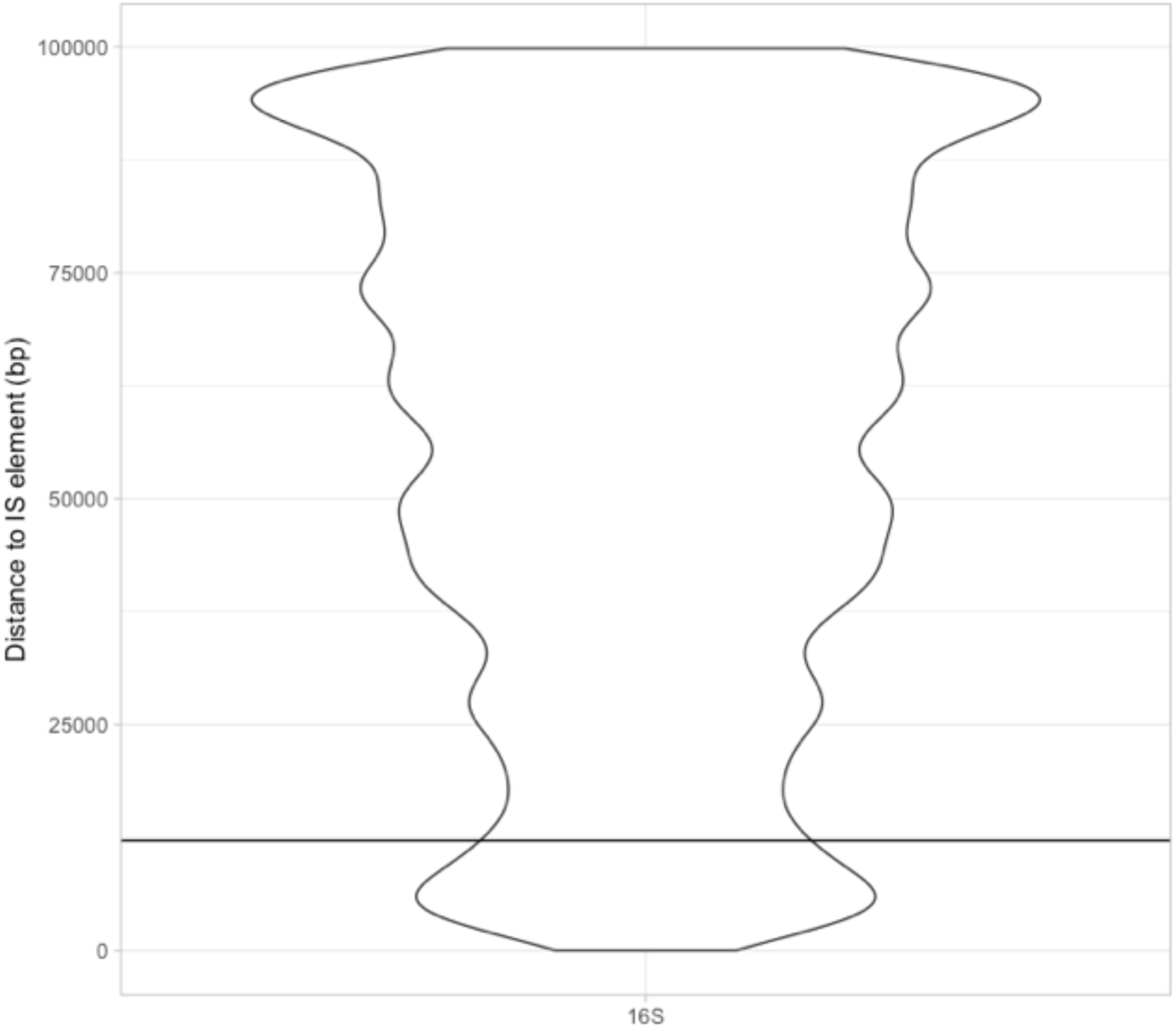
Violin plot of distances between all 16S rRNA genes and the closest IS element. Only 16S:IS associations within 100 kbp were plotted here.

### Supplementary Note 3: Database biases

In the RefSeq complete genomes, bacteria belonging to the Enterobacterales order make up the biggest order in the database and represent 20.42% of entries (Supplementary Fig. 6). Likewise, specialized functional gene databases, such as CARD and ISfinder, can be assumed to be biased towards certain taxonomic groups. Together, these biases will likely bias the analyses presented here synergistically by the combined biases the RefSeq, CARD, and ISfinder databases.

**Supplementary Fig. 6.**
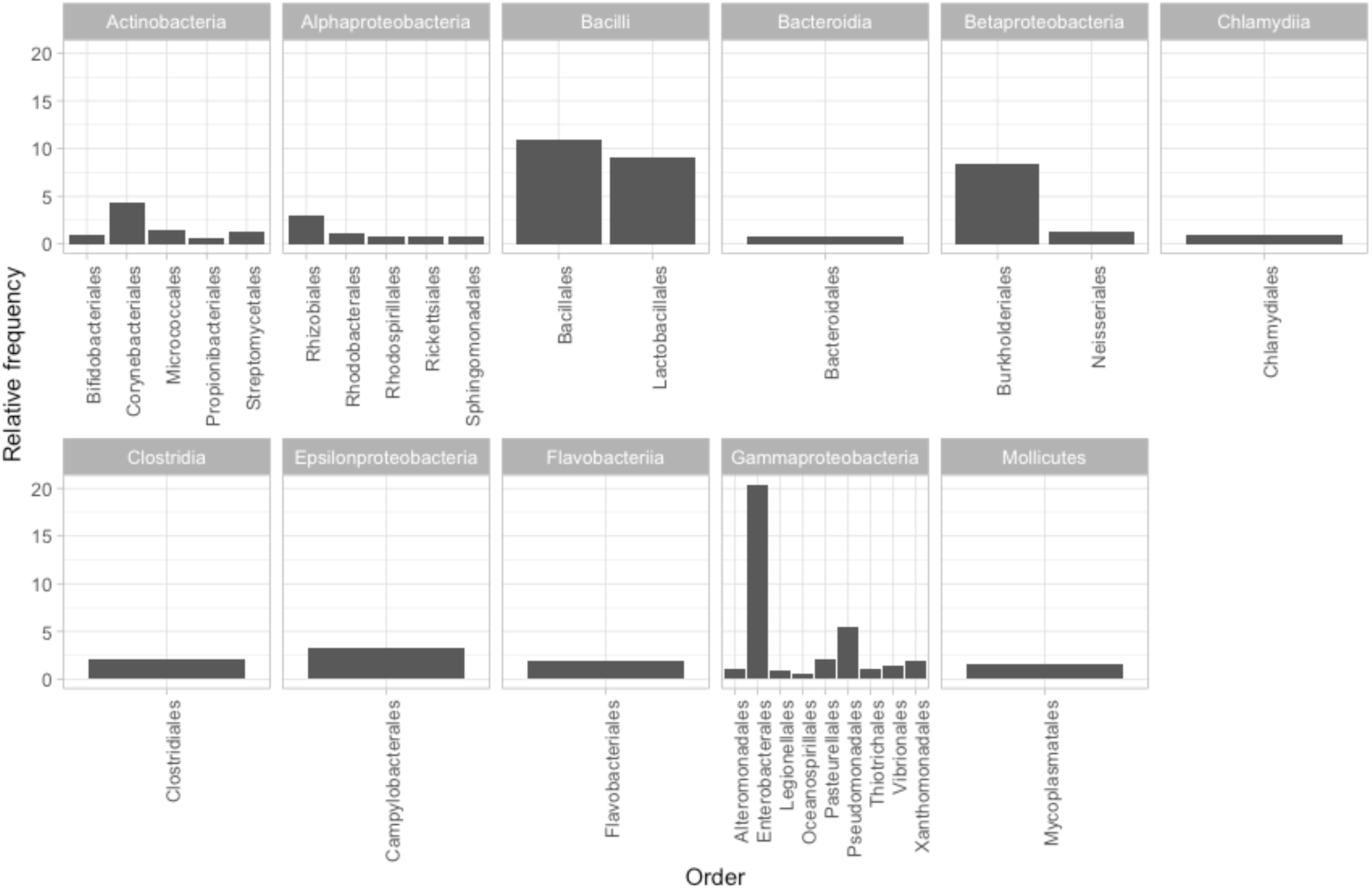
Frequencies (%) of RefSeq genomes by order taxonomic level. Only orders making up at least 0.5% of the total abundance are shown. Orders are organized by phylum.

Proteins in the CARD database are representatives from single strains of bacteria. Comparing relative abundances of represented genera in the CARD protein homology database with the relative abundances of the same genera in the RefSeq complete genome database shows that the two databases do not have an equal distribution of genera (Supplementary Fig. 7).

**Supplementary Fig. 7.**
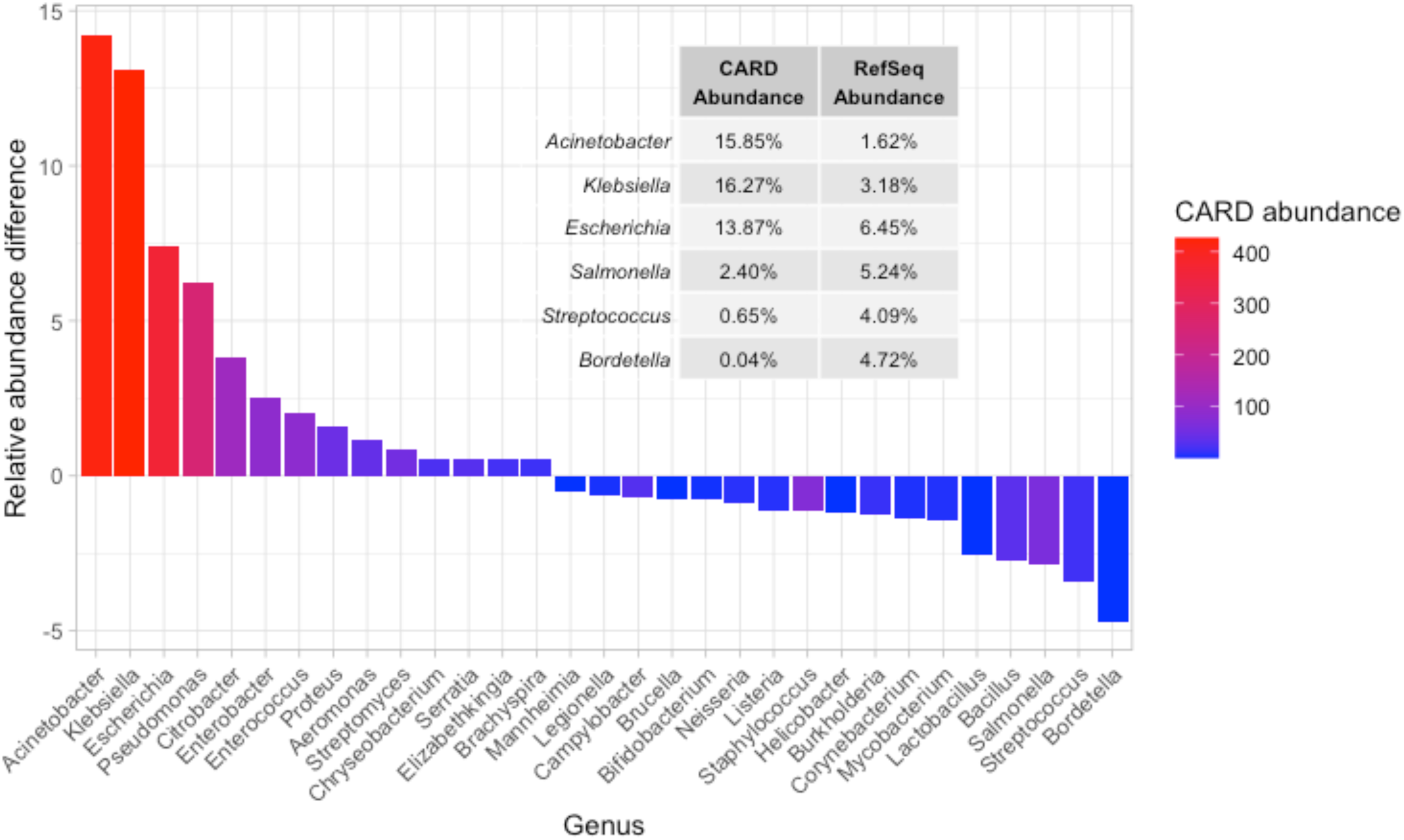
Differences in relative abundance in representation of genera in CARD and RefSeq complete genomes databases. Positive difference indicates higher abundance in CARD than in RefSeq and vice versa. Only genera present in both databases and with difference in relative abundance of at least 0.5% are shown.

The 6 most extreme cases with biggest difference in relative abundance between CARD and RefSeq databases are highlighted in a table within Supplementary Fig. 7. Putative ARGs from *Acinetobacter* strains represent 15.6% of data in CARD, while *Acinetobacter* only make up 1.6% of entries in RefSeq complete genomes. Conversely, *Bordetella* genomes constitute 4.7% of RefSeq complete genomes, but are only represented by 0.04% of the CARD proteins. *Streptococcus* is likewise less abundant in CARD. All of the top 6 most different genera are all of potential clinical relevance and there is no apparent logical reason why one of these genera should have fewer or more inherent or acquired ARGs than the other. Considering the absolute abundance of the CARD genera, it seems likely that the differences in relative abundance between CARD and RefSeq is largely due to biased representation in CARD proteins. ARG prediction in underrepresented genera such as *Bordetella* and *Streptococcus* is therefore likely to be less accurate and encompassing than in e.g. *Acinetobater*, *Klebsiella*, and *Escherichia*.

### Supplementary Note 4: Ameliorating database biases by clustering

Most publicly available sequence databases are biased in entries towards organisms that have gathered the most research interest, usually human-associated bacteria such as enterobacteria. It is therefore assumed that both CARD and RefSeq databases are heavily biased, but not towards the same genera. These biases will, naturally, affect analyses performed in this study. However, as one of the most curated and widely used ARG databases, CARD is the only obvious choice for this study. Likewise, RefSeq complete genomes comprise a large and well-curated database of publicly available genomes and is the natural choice for this study. Both RefSeq and CARD databases are generally biased towards the Enterobacteriales order but RefSeq is more skewed towards *Bordetella*, *Streptococcus*, and *Salmonella* than CARD. On the other hand, CARD entries are overrepresented by *Acinetobacter*, *Klebsiella*, and *Escherichia* compared to RefSeq (Supplementary Figs. 7,8a).

**Supplementary Fig. 8:**
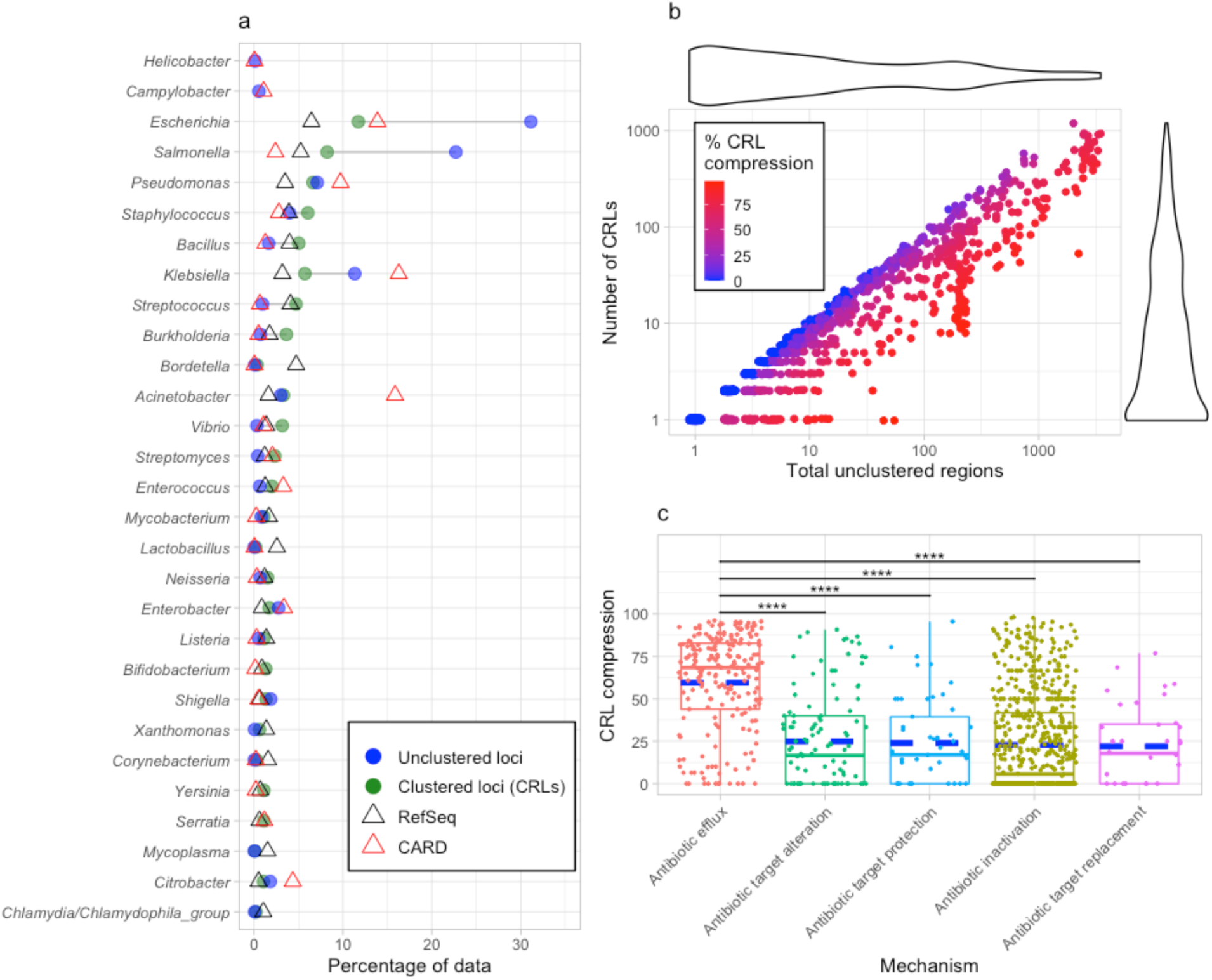
Effect of clustering ARG loci to CRLs. a, Database content of most abundant genera and the effect of clustering to CRLs. Clustered loci (CRLs; green circles), divided into the parent genus, generally come closer to the relative abundance of the given genus in the RefSeq complete genomes database (black triangles), than the unclustered, redundant ARG loci (blue circles). The relative abundance of genera from the CARD database are also shown (red triangles). Only genera that make up at least 1% of the total dataset of either unclustered loci, CRLs, or RefSeq complete genomes are shown. The displayed genera constitute 92.68%, 78.43%, 61.64%, and 81.52% of the total datasets for unclustered loci, CRLs, RefSeq, and CARD databases, respectively. b, Compression effect of clustering ARG loci to CRLs. Each dot represents a single ARO category where its position on the x-axis indicates the number of identified unclustered loci and its position on the y-axis is the number of CRLs produced from clustering. The colour of the points shows the compression rate of clustering in percentage, calculated by dividing number of CRLs with the number of unclustered loci. Red points indicate that there are many, almost identical, ARG loci for a given ARO category, resulting in a low number of CRLs relative to the number of unclustered loci. Vice versa, blue points indicate that most of the unclustered loci are unique and clustering results in a number of CRLs that is close to the number of total unclustered loci. Violin plots are shown for both axes on the outside of the plot. Note that the position of points have been jittered very slightly to improve visualization. c, The CRL compression rate per major resistance mechanism (the very low abundance ‘Reduced permeability to antibiotic’ category is not shown). Each jittered point is a unique ARO CRL. Boxplot beneath CRL points show interquartile range and median as solid horizontal line. The dashed blue line indicates the mean. Whiskers extend to 1.5 * of the interquartile ranges and outliers of this range are not highlight. Difference in means of CRL compression per resistance mechanism were tested with Mann-Whitney U-test with FDR correction for multiple testing. Only significant differences are shown (**** P < 0.0001).

In order to minimize the impact of oversampling of almost identical genomes from RefSeq (e.g. *E. coli* substrains), ARG loci were clustered with USEARCH to 99% sequence similarity over at least 90% of the region length (length of ARG + 12,170 bp in both directions). 176,688 loci with ARGs passed all filters and were clustered to 53,895 CRLs that represent 1,176 individual CARD ARO categories out of a total of 2,617 in the CARD protein homolog database (Supplementary Table 2). The missing AROs are located in bacteria that do not yet have completed genomes or are so similar to one of the 1,176 identified AROs that they were not included, since only the best ARO match per query protein was considered. Indeed, removing the initial 1,176 AROs from the CARD database and performing the analyses again resulted in 336 AROs that were not included in the main analysis (results not shown).

**Supplementary Table 2.**
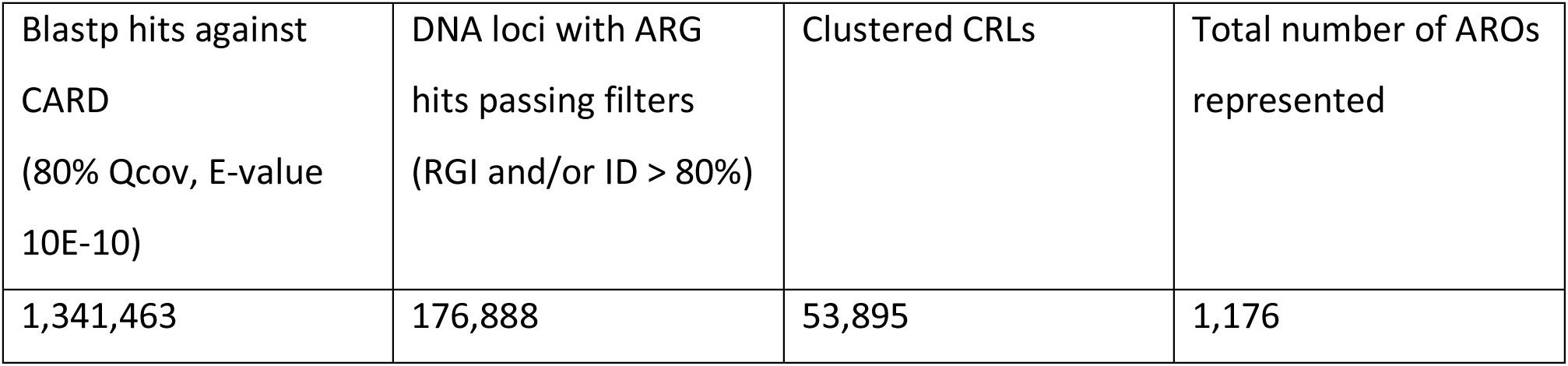
Of the 176,888 loci passing filters, 115,268 pass the RGI bitscore cutoff while 61,620 pass the 80% ID filter. All hits have at least 80% query coverage against the CARD ARGs.

| Blastp hits against<br>CARD<br>(80% Qcov, E-value<br>10E-10) | DNA loci with ARG<br>hits passing filters<br>(RGI and/or ID > 80%) | Clustered CRLs | Total number of AROs<br>represented |
| --- | --- | --- | --- |
| 1,341,463 | 176,888 | 53,895 | 1,176 |

Genetic loci with ARGs are summarized based on the genus of the strain they occur in (Supplementary Fig. 8a). The relative abundance of each genus with ARGs are compared with the relative abundance in the RefSeq complete genome database. The difference between relative abundances per genus (given in percent of total database size) are summarized, resulting in a Euclidean distance of 30.89 (373 genera) between unclustered loci and RefSeq. When clustering these loci to CRLs, the Euclidean distance of relative genera abundances compared to RefSeq abundance is reduced to 10.26 (370 genera), which shows that clustering reduces effect of oversampling of e.g. almost clonal *E. coli* strains. The most abundant genera, which make up at least 1% of either unclustered loci, CRLs, or RefSeq, are shown in Supplementary Fig. 8a. Almost all of the 29 genera shown are associated with either human pathogens or other anthropogenic activity (*Lactobacillus*, *Bifidobacterium*), showing that bacteria in these ecological niches are overrepresented in the both the RefSeq and the CARD database. The CARD database is even more biased towards known pathogens including *Escherichia*, *Pseudomonas*, *Klebsiella*, *Acinetobacter*, and a few others (Supplementary Fig. 8a). This bias is sure to have a major effect on ARG prediction and leads to high ARG estimates in these genera compared to others (e.g. environmental bacteria). Clustering nearly identical ARG loci to CRLs definitely helps to smoothen this skew but it cannot completely even out the biases discussed here. Furthermore, clustering to CRLs reduces the overemphasis on the human-associated genera shown in Supplementary Fig. 8a from 92.68% of the total ARG loci to 78.43%, which slightly improves representation of other genera. These database biases are not surprising, since many researchers and clinicians are interested in human pathogens or closely related bacteria that can develop and transfer antimicrobial resistance. Therefore, we accept these biases in the present study but are aware that environmental bacteria and their potential resistance genes are underrepresented here.

Not all ARO categories are compressed equally in relative abundance by clustering loci to CRLs (Supplementary Fig. 8b,c). Some AROs are represented only by completely unique DNA loci in the unclustered dataset, resulting in a number of CRLs that is the same as the number of unclustered loci (low compression rate). On the other hand, other AROs are represented by a large number of nearly identical DNA loci which either stem from biased oversampling of e.g. almost clonal *E. coli* or by clonal expansion of one or more DNA loci by HGT (high compression rate).

Generally, loci with efflux pump resistance determinants have a significantly higher average CRL compression rate than the other functional categories (Figure 1, Supplementary Fig. 8c; P < 0.0001), which means that there are many almost clonal DNA loci with efflux pumps in the RefSeq complete genome database. The other general functional categories do not have a significantly different CRL compression rate, signifying that AROs belonging to these categories are located in more diverse DNA loci than efflux pumps are.

**Supplementary Fig. 9.**
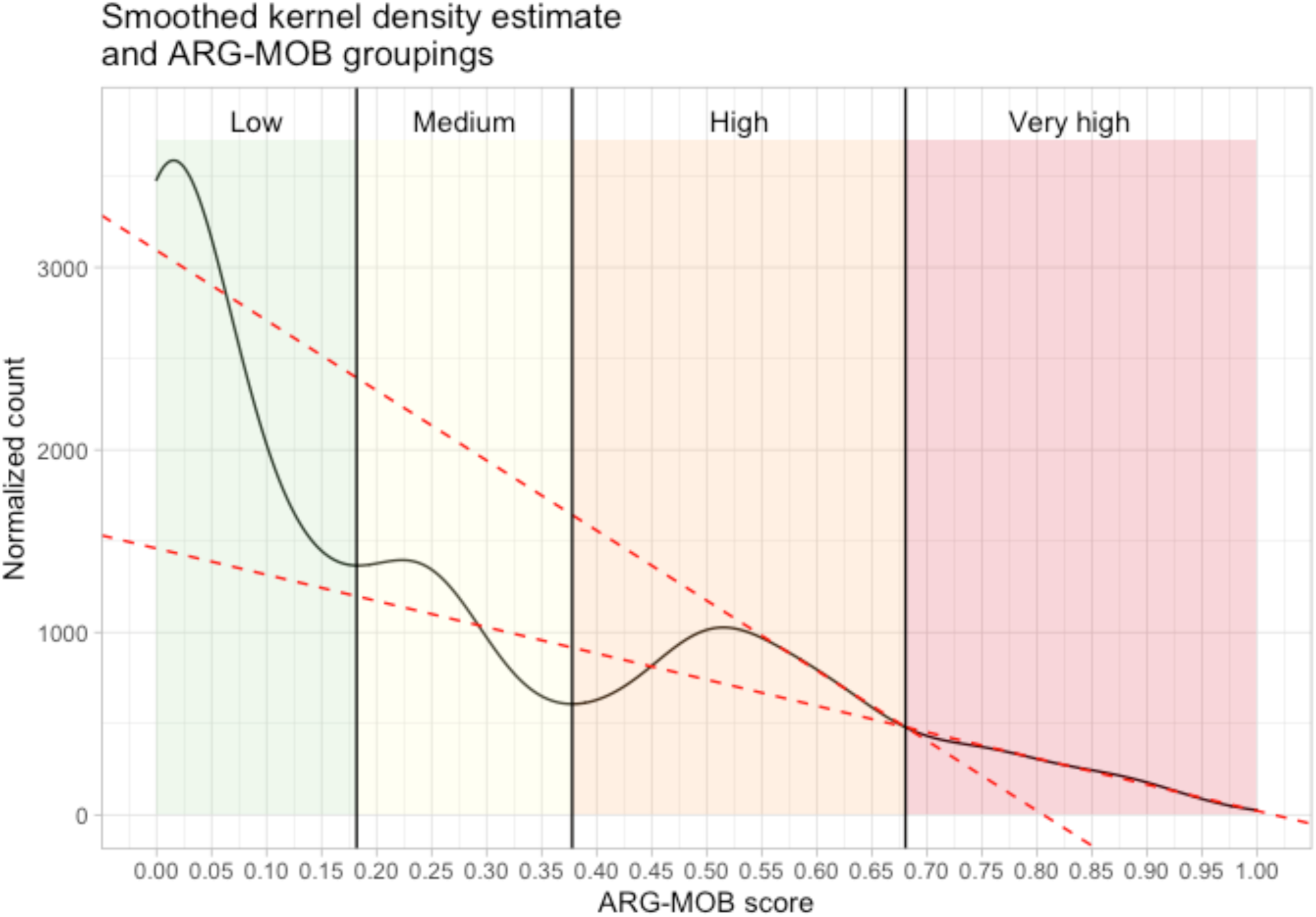
Smoothed kernel density estimates of all AROs and their ARG-MOB values. ARG-MOB values defining the categories were computed by identifying local minima in R. The High/Very high intersection is not found by this method. Instead, linear models were fitted to High interval 0.55-0.65 and Very high interval 0.7-1. These intervals are approximately linear. The intersection between the two models (0.685) is used as the High-Very high limit.

### Supplementary Note 5: IS element families in major mechanisms

The IS elements occurring in the proximity of putative ARGs were classified by their IS families and tested for significantly different relative abundances between resistance mechanisms (Supplementary Fig. 10). Among the 17 most abundant IS families, IS*1*, IS*110*, IS*1595*, IS*1380*, IS*200*/IS*605*, IS*21*, IS*3*, IS*30*, IS*4*, IS*5*, IS*6*, IS*66*, IS*91*, ISL*3*, and Tn*3* occur with significantly different frequencies in AI and AE loci, with lower median frequencies in AE. These families are therefore more active in decontextualizing putative AI ARGs than AE, accompanying the observation that AI is generally more mobilized than AE. CRLs of the ATR mechanism are significantly less associated with a lack of IS elements, which in turn means that there are more CRLs of this mechanism with IS elements in proximity. This is also shown in Figure 3 and Supplementary Fig. 11 where it is apparent that this mechanism is very often found in association with IS elements and often on plasmids.

**Supplementary Fig. 10.**
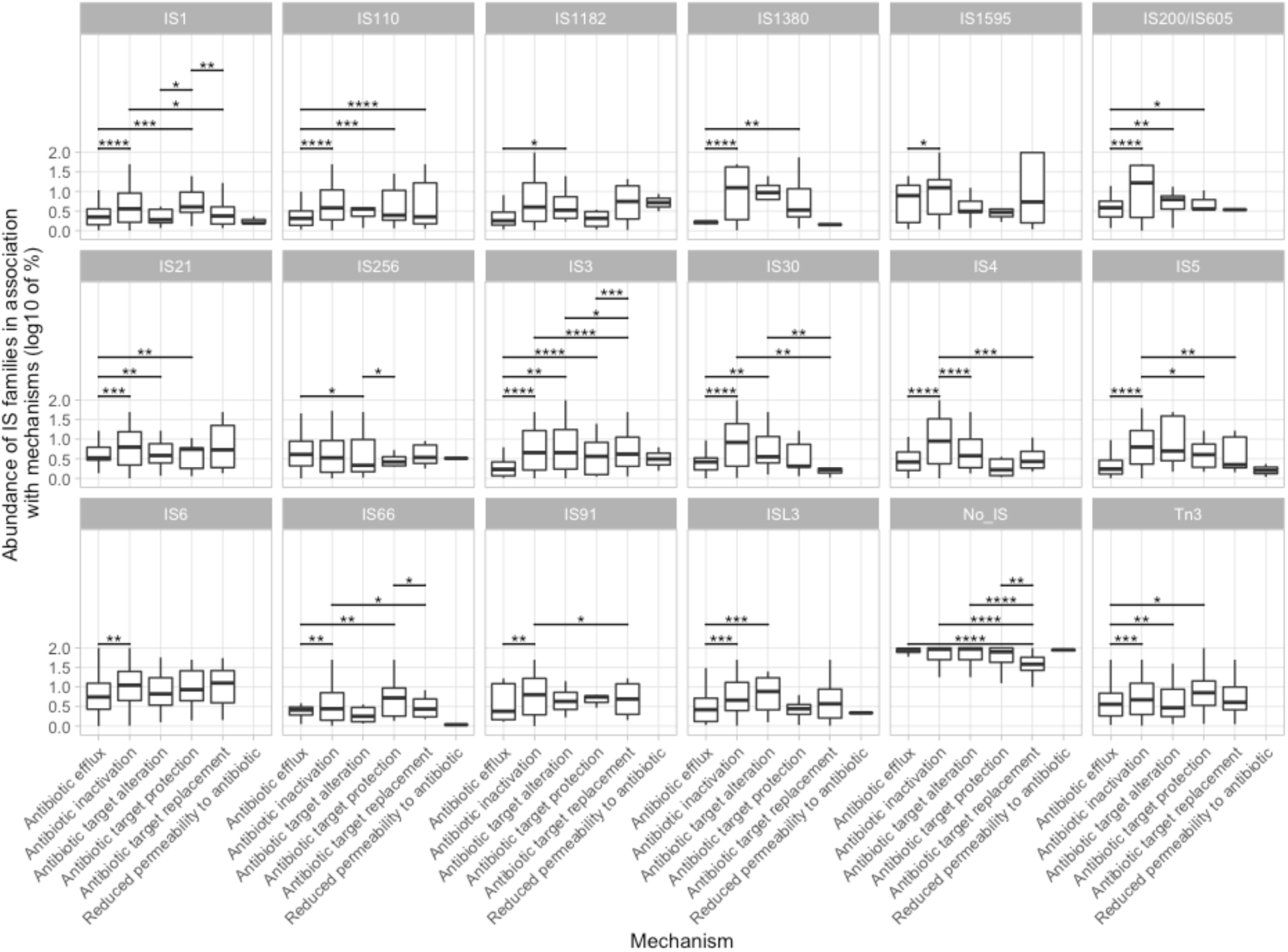
Relative abundance of 17 IS families and the lack of IS elements (No_IS) in ARO mechanisms. Only IS families with significant differences in mean value between mechanisms are shown. Log10 of relative abundance on Y-axes is derived from the frequency of IS families per mechanism category. Boxes indicate first and third quartiles (25% and 75% of data) and horizontal line in box shows the median. Whiskers extend to 1.5 * of the interquartile ranges. Above boxplots, bars indicate significant differences in mean between mechanisms (Mann-Whitney test with FDR correction). Only significant differences are displayed (****: p <= 0.0001).

### Supplementary Note 6: Antibiotic efflux ARGs are more loosely associated with IS elements

Within the 12,170 bp investigated in both directions of identified ARGs, the distance to nearest IS element might be thought to be an indicator of how “tightly” associated a given ARG is with an IS element. Likely, there are some false positive associations between ARGs and IS elements found in the extremes of the 12,170 bp maximum distance. The mean distance between ARGs and IS elements per ARO is shown on the Y-axes in Supplementary Fig. 11a (mean distance per ARO is calculated from unique CRLs). For AE, the bulk of the AROs have low IS ratios and a mean distance to closest IS elements of just over 5000 bp. This is significantly different from 4 of the 5 other major mechanisms (Supplementary Fig. 11b), with the exception for the very low abundance Reduced permeability to antibiotic mechanism. The AE AROs that do have higher IS ratios also have shorter distances to closest IS elements, which is comparable to AROs of other mechanisms that are also high mobilized. For these mobilized AROs, the mean distance between ARGs and IS elements is closer to 2,500 bp.

**Supplementary Fig. 11:**
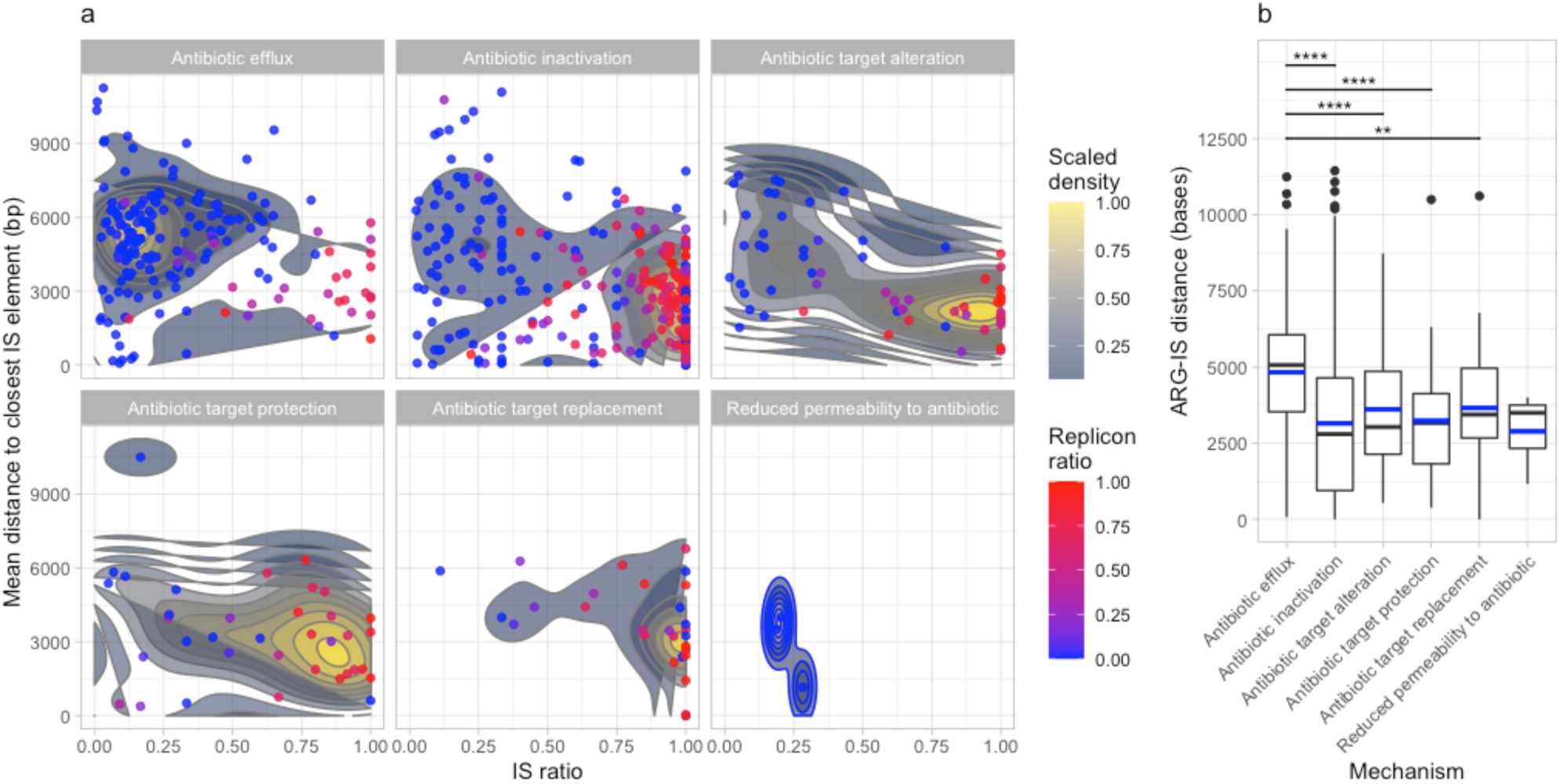
Distance between ARGs and closest IS element. a, Density plots of IS ratio against the mean distance in bases to the nearest IS element in both directions. Each point represents a unique ARO category. Plots are divided into the individual mechanisms and are colored according to the replicon ratio, where a high ratio (red) indicates that an ARO is more often found on plasmids and a low ratio (blue) indicates that an ARO is more on chromosomes. Density estimates are calculated with two-dimensional kernel density estimation, as implemented in the stat_density_2d function under the ggplot R package. b, Boxplot of median distance (bases) between ARGs and closest IS elements. Mean is shown with blue dashed lines. Boxes indicate first and third quartiles (25% and 75% of data) and horizontal line in box shows the median. Whiskers extend to 1.5 * of the interquartile ranges. Outliers are shown as black dots. Above boxplots, bars indicate significant differences in mean between mechanisms (Mann-Whitney test with FDR correction). Only significant differences are displayed (**: p <= 0.01, ****: p <= 0.0001).

### Supplementary Note 7: Taxonomic investigation

Within Proteobacteria, *Enterobacteriaceae* have a higher median IS ratio than *Campylobacteriaceae,* and *Burkholderiaceae* but lower than *Aeromonadaceae*, *Pasterurellaceae* and *Morganellaceae* (Supplementary Fig. 12). However, CRLs in *Enterobacteriaceae* are more often found on plasmids than for *Campylobacteraceae*, *Moraxellaceae*, *Morganellaceae*, *Neisseriaceae*, *Pasteurellaceae*, *Pseudomonadaceae*, and *Burkholderiaceae* (Supplementary Fig. 13; MWU test; P < 0.01), showing that many putative ARGs in *Enterobacteriaceae* are highly mobilized by both IS elements and plasmids.

Within *Enterobacteriaceae*, many putative ARG loci have been mobilized both by IS elements and plasmids, especially within genera *Shigella*, *Escherichia*, *Salmonella*, *Klebsiella*, *Enterobacter*, and to a lesser degree *Citrobacter* (Supplementary Fig. 12c). Other genera in *Enterobacteriaceae* show lower mean mobilization degrees (Significance values in Supplementary Table 3). *Enterobacteriaceae* genera with highly mobilized ARGs all have members of significant importance to human health and persistent fixation of mobilized ARGs are likely a consequence of human interference with pathogenic bacteria^3^.

Phyla whose members are more associated with the environment, such as Actinobacteria and Bacteriodetes, have lower median IS ratios than Proteobacteria (MWU test; P < 0.01). However, within e.g. Firmicutes, whose IS ratio is not different from Proteobacteria, some families are also associated with human activities such as *Enterococcaceae* and *Staphylococcaceae*. These also harbour highly mobilized ARGs, while environmentally associated Firmicutes, such as *Bacillaceae*, barely have ARGs mobilized by either IS elements or plasmids (Supplementary Fig. 14). This exemplifies how homologs of ARGs can be found in both environmental and clinically relevant genera, but that they have been decontextualized more in the latter.

**Supplementary Fig. 12:.**
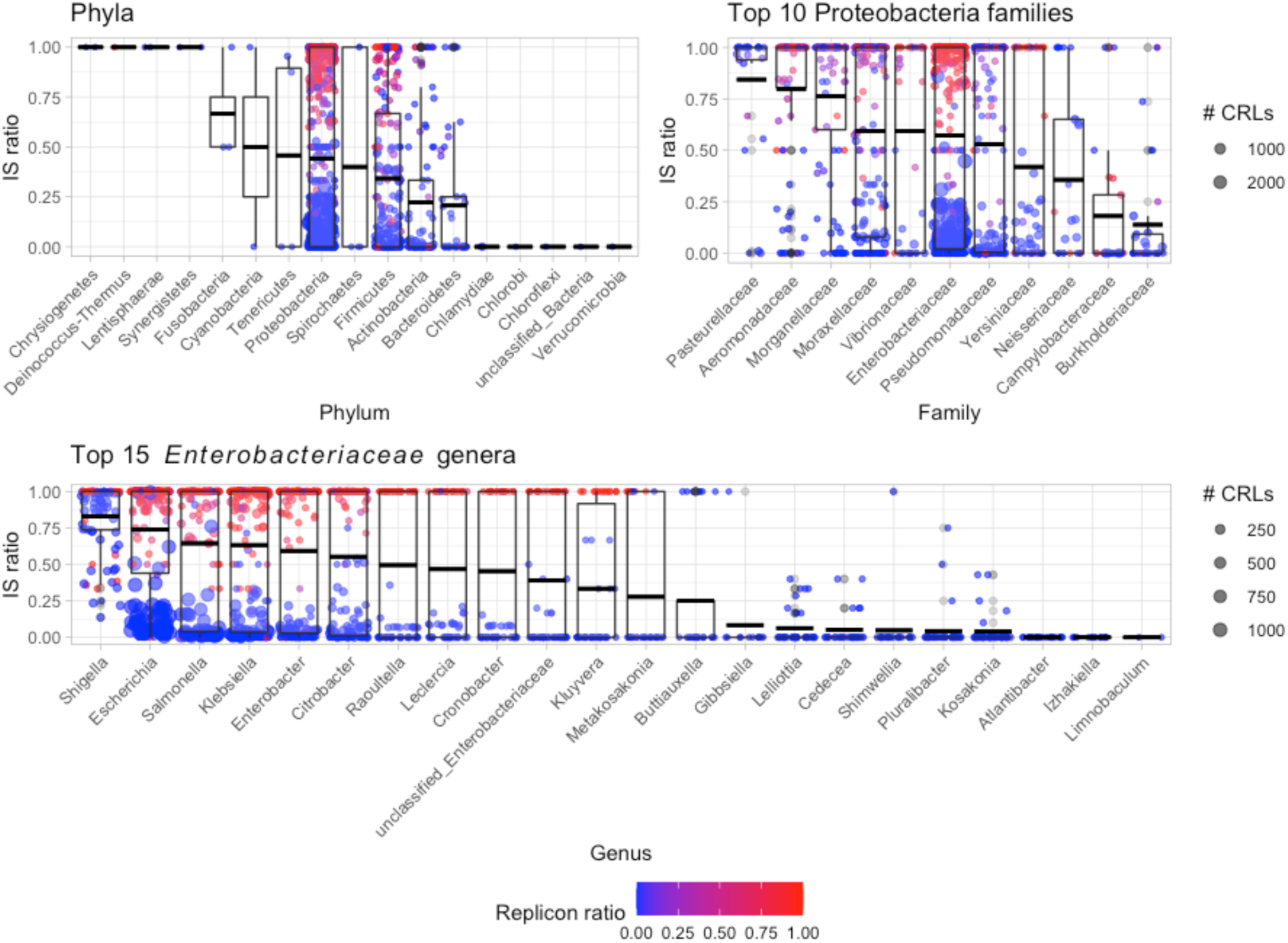
Taxonomic distribution of ARO categories. Boxplots and dots show IS ratio per taxonomic group. The size of the points indicates the number of unique CRLs in a given ARO, while the colour is the replicon ratio with highest (red) indicating more plasmid than chromosome placement of CRLs. Focus is placed on the Proteobacteria for plotting of families and genera. Boxes indicate first and third quartiles (25% and 75% of data) and horizontal line in box shows the median. Whiskers extend to 1.5 * of the interquartile ranges.

**Supplementary Fig. 13.**
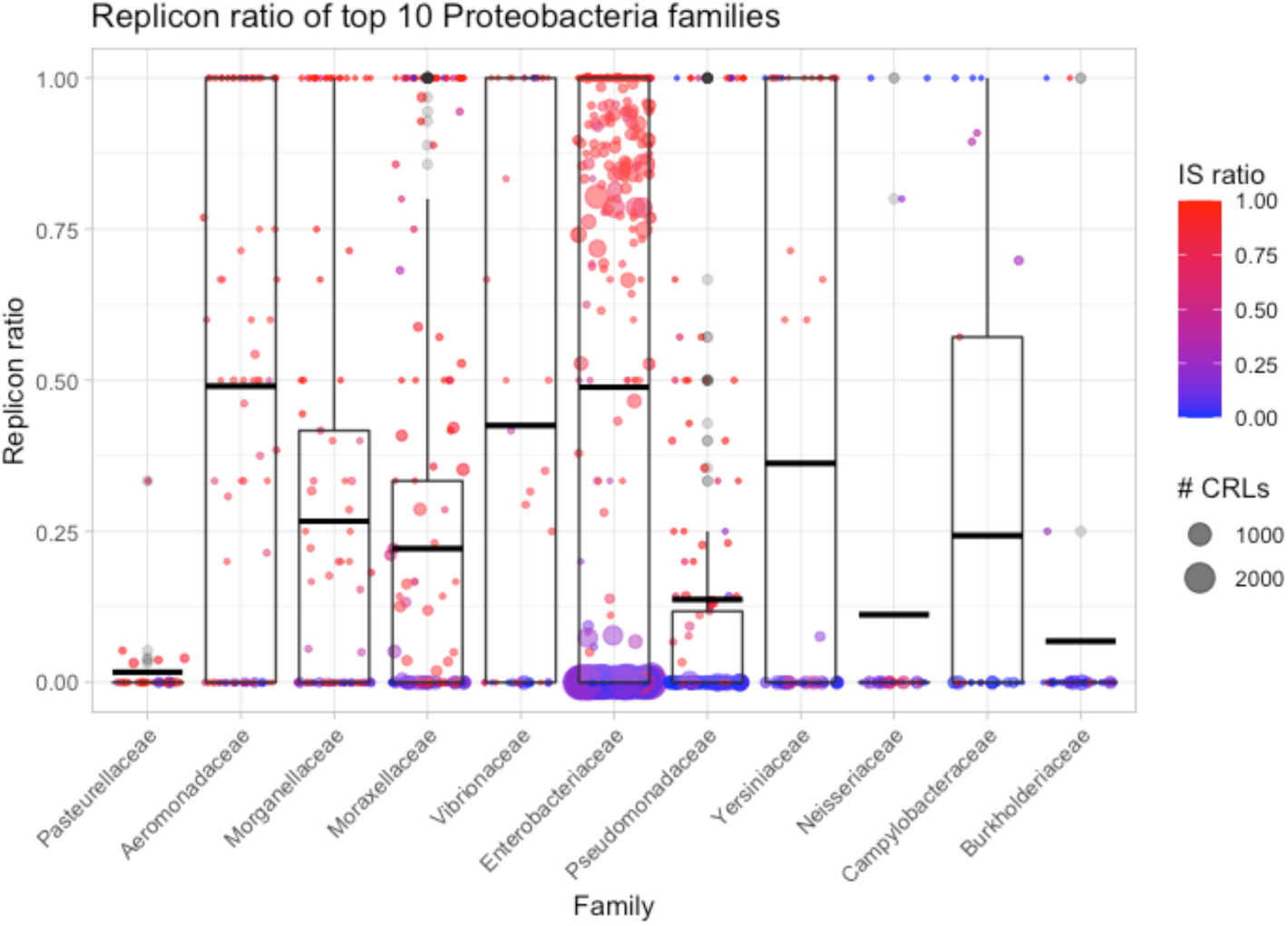
Replicon ratio of the top 10 most abundant Proteobacterial families is shown as boxplots. The colour of the CRL circles indicate IS ratio rather than the Replicon ratio shown in Supplementary Fig. 12.

**Supplementary Fig. 14.**
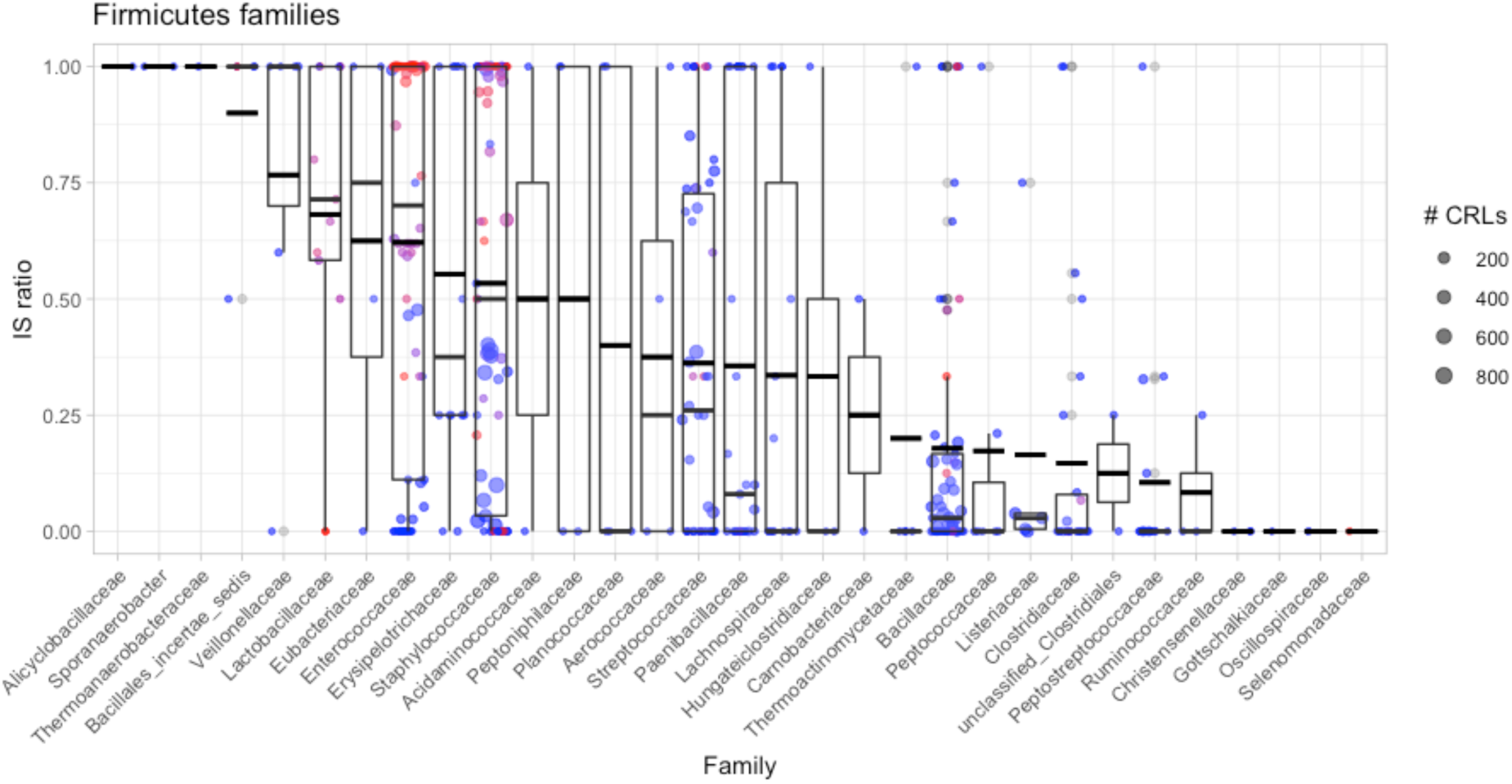
Taxonomic distribution of ARO categories within the Firmicutes phylum. Boxplots and dots show IS ratio per taxonomic group. The size of the points indicates the number of unique CRLs in a given ARO, while the colour is the replicon ratio with highest (red) indicating more plasmid than chromosome placement of CRLs Boxes indicate first and third quartiles (25% and 75% of data) and horizontal line in box shows the median. Whiskers extend to 1.5 * of the interquartile ranges.

**Supplementary Table 3.** Pairwise Mann-Whitney tests for IS and Replicon ratios in *Enterobacteriaceae* with FDR correction for multiple testing. Only significant comparisons are listed here. Significance levels are *: p < 0.05; **: p < 0.01; ***: p < 0.001; ****: p < 0.0001.

| Genus 1 | Genus 2 | IS p | IS sign. | Replicon p | Replicon sign. |
| --- | --- | --- | --- | --- | --- |
| *Cedecea* | *Citrobacter* | 4.70E-08 | **** | 8.10E-06 | **** |
| *Cedecea* | *Cronobacter* | 3.00E-04 | *** | 0.00026 | *** |
| *Cedecea* | *Enterobacter* | 6.00E-09 | **** | 2.50E-07 | **** |
| *Cedecea* | *Escherichia* | 1.70E-14 | **** | 2.20E-09 | **** |
| *Cedecea* | *Klebsiella* | 4.60E-10 | **** | 2.30E-08 | **** |
| *Cedecea* | *Kluyvera* | 0.027 | * | 0.0078 | ** |
| *Cedecea* | *Leclercia* | 3.60E-05 | **** | 0.00021 | *** |
| *Cedecea* | *Raoultella* | 3.60E-05 | **** | 2.70E-05 | **** |
| *Cedecea* | *Salmonella* | 5.20E-12 | **** | 8.20E-08 | **** |
| *Cedecea* | *Shigella* | 5.30E-14 | **** | 0.00027 | *** |
| *Cedecea* | *unclassified_Enterobacteriaceae* | 0.0048 | ** | 0.00084 | *** |
| *Citrobacter* | *Escherichia* | 0.00023 | *** | 0.0023 | ** |
| *Citrobacter* | *Kluyvera* | 0.0012 | ** | 0.045 | * |
| *Citrobacter* | *Kosakonia* | 1.50E-10 | **** | 4.20E-07 | **** |
| *Citrobacter* | *Lelliottia* | 1.60E-09 | **** | 1.60E-07 | **** |
| *Citrobacter* | *Pluralibacter* | 2.40E-11 | **** | 4.20E-07 | **** |
| *Citrobacter* | *Shigella* | 0.0098 | ** | NA | NA |
| *Citrobacter* | *unclassified_Enterobacteriaceae* | 0.0071 | ** | NA | NA |
| *Cronobacter* | *Escherichia* | 0.00082 | *** | NA | NA |
| *Cronobacter* | *Kosakonia* | 1.80E-05 | **** | 4.20E-05 | **** |
| *Cronobacter* | *Lelliottia* | 0.00022 | *** | 2.60E-05 | **** |
| *Cronobacter* | *Pluralibacter* | 2.30E-06 | **** | 4.20E-05 | **** |
| *Cronobacter* | *Salmonella* | 0.023 | * | NA | NA |
| *Cronobacter* | *Shigella* | 0.006 | ** | NA | NA |
| *Enterobacter* | *Escherichia* | 0.0037 | ** | NA | NA |
| *Enterobacter* | *Kluyvera* | 0.00019 | *** | 0.0031 | ** |
| *Enterobacter* | *Kosakonia* | 1.30E-11 | **** | 5.70E-09 | **** |
| *Enterobacter* | *Lelliottia* | 1.30E-10 | **** | 1.70E-09 | **** |
| *Enterobacter* | *Pluralibacter* | 1.50E-12 | **** | 5.70E-09 | **** |
| *Enterobacter* | *unclassified_Enterobacteriaceae* | 0.0015 | ** | 0.032 | * |
| *Escherichia* | *Klebsiella* | 0.0041 | ** | NA | NA |
| *Escherichia* | *Kluyvera* | 3.40E-08 | **** | 0.00022 | *** |
| *Escherichia* | *Kosakonia* | 4.90E-18 | **** | 3.70E-11 | **** |
| *Escherichia* | *Leclercia* | 0.0012 | ** | NA | NA |
| *Escherichia* | *Lelliottia* | 8.50E-18 | **** | 1.60E-11 | **** |
| *Escherichia* | *Pluralibacter* | 2.70E-18 | **** | 3.70E-11 | **** |
| *Escherichia* | *Raoultella* | 0.0018 | ** | NA | NA |
| *Escherichia* | *unclassified_Enterobacteriaceae* | 2.40E-07 | **** | 0.0029 | ** |
| *Klebsiella* | *Kluyvera* | 6.80E-05 | **** | 0.00087 | *** |
| *Klebsiella* | *Kosakonia* | 5.80E-13 | **** | 4.40E-10 | **** |
| *Klebsiella* | *Lelliottia* | 2.30E-12 | **** | 1.10E-10 | **** |
| *Klebsiella* | *Pluralibacter* | 1.10E-13 | **** | 4.40E-10 | **** |
| *Klebsiella* | *unclassified_Enterobacteriaceae* | 0.00053 | *** | 0.0099 | ** |
| *Kluyvera* | *Kosakonia* | 0.0058 | ** | 0.0025 | ** |
| *Kluyvera* | *Leclercia* | 0.034 | * | NA | NA |
| *Kluyvera* | *Lelliottia* | 0.029 | * | 0.0017 | ** |
| *Kluyvera* | *Pluralibacter* | 0.0017 | ** | 0.0025 | ** |
| *Kluyvera* | *Raoultella* | 0.028 | * | 0.032 | * |
| *Kluyvera* | *Salmonella* | 2.50E-06 | **** | 0.0034 | ** |
| *Kluyvera* | *Shigella* | 1.20E-06 | **** | NA | NA |
| *Kosakonia* | *Leclercia* | 1.00E-06 | **** | 3.30E-05 | **** |
| *Kosakonia* | *Raoultella* | 1.20E-06 | **** | 2.50E-06 | **** |
| *Kosakonia* | *Salmonella* | 2.40E-15 | **** | 1.70E-09 | **** |
| *Kosakonia* | *Shigella* | 1.10E-16 | **** | 4.20E-05 | **** |
| *Kosakonia* | *unclassified_Enterobacteriaceae* | 0.00061 | *** | 0.00016 | *** |
| *Leclercia* | *Lelliottia* | 1.70E-05 | **** | 1.80E-05 | **** |
| *Leclercia* | *Pluralibacter* | 1.10E-07 | **** | 3.30E-05 | **** |
| *Leclercia* | *Shigella* | 0.0041 | ** | NA | NA |
| *Lelliottia* | *Raoultella* | 1.50E-05 | **** | 1.10E-06 | **** |
| *Lelliottia* | *Salmonella* | 2.30E-14 | **** | 4.60E-10 | **** |
| *Lelliottia* | *Shigella* | 2.10E-17 | **** | 2.60E-05 | **** |
| *Lelliottia* | *unclassified_Enterobacteriaceae* | 0.0037 | ** | 8.90E-05 | **** |
| *Pluralibacter* | *Raoultella* | 1.60E-07 | **** | 2.50E-06 | **** |
| *Pluralibacter* | *Salmonella* | 3.00E-16 | **** | 1.70E-09 | **** |
| *Pluralibacter* | *Shigella* | 2.00E-16 | **** | 4.20E-05 | **** |
| *Pluralibacter* | *unclassified_Enterobacteriaceae* | 0.00015 | *** | 0.00016 | *** |
| *Raoultella* | *Shigella* | 0.02 | * | NA | NA |
| *Salmonella* | *unclassified_Enterobacteriaceae* | 2.40E-05 | **** | 0.043 | * |
| *Shigella* | *unclassified_Enterobacteriaceae* | 2.80E-05 | **** | NA | NA |
| *Citrobacter* | *Klebsiella* | NA | NA | 0.011 | * |
| *Enterobacter* | *Shigella* | NA | NA | 0.011 | * |
| *Escherichia* | *Salmonella* | NA | NA | 0.036 | * |
| *Escherichia* | *Shigella* | NA | NA | 0.00026 | *** |
| *Klebsiella* | *Shigella* | NA | NA | 0.0016 | ** |
| *Salmonella* | *Shigella* | NA | NA | 0.016 | * |

### Supplementary Note 8: Supplementary information to integron analyses

**Supplementary Fig. 15:**
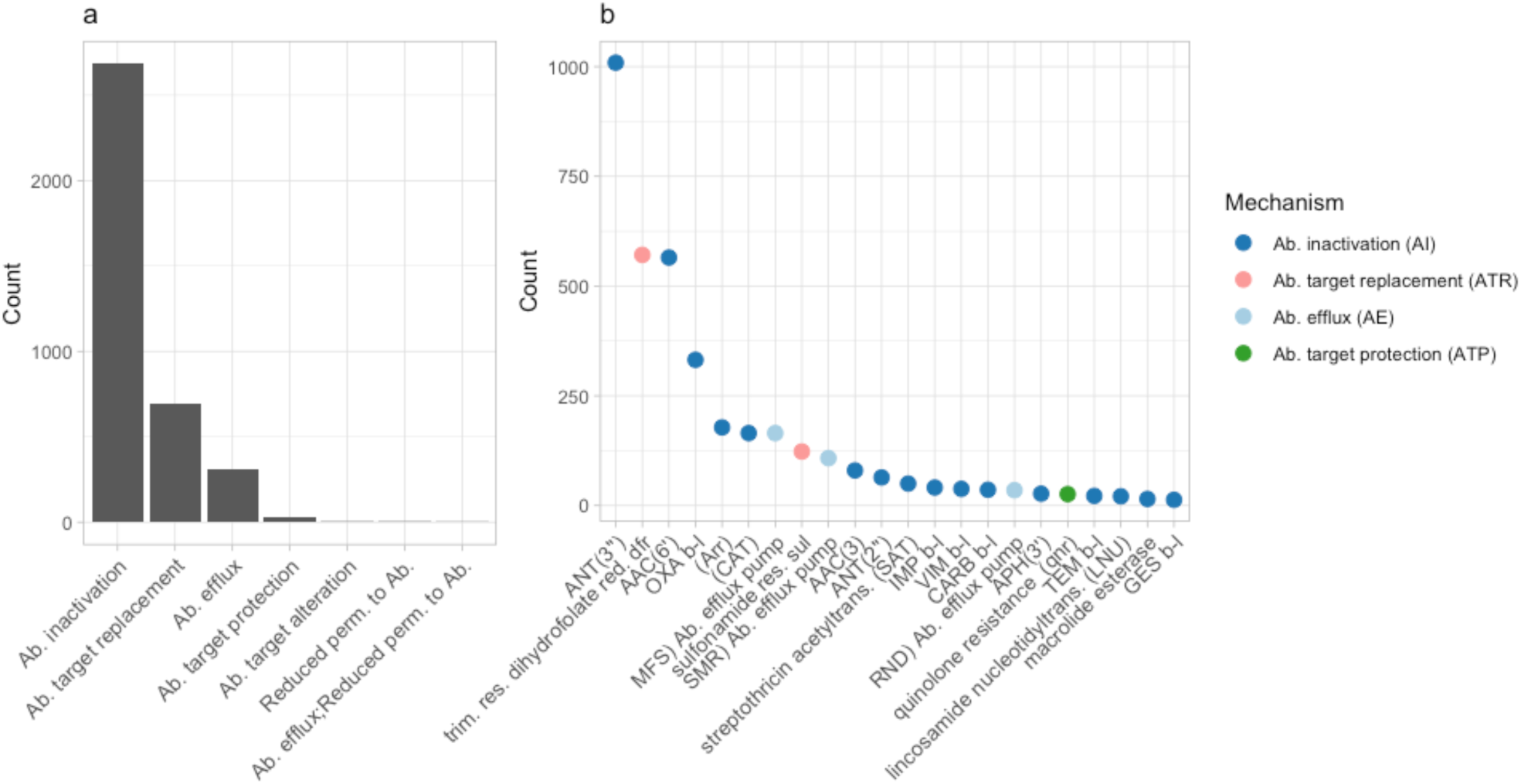
ARGs associated with integrons. a, Major mechanisms associated with integrons. The count is in unique CRLs belonging to each mechanism. b, CRLs where the identified ARG is inserted in an integron, summarized by submechanism category. Each point can include multiple ARO categories. Only submechanisms with more than 10 CRLs are shown.

The 5 ANT AROs (*aadA*, *aadA2*, *aadA5*, *ANT(3’’)-IIa*, and *ANT(2’’)-Ia)* shown in Supplementary Table 4 all display integron associations in at least 54.24% of their total CRL occurrences. Similarly, the aminoglycoside acetyltransferase *AAC(6’)* AROs (*AAC(6’)-Ib-cr*, *AAC(6’)-Ib10*, *AAC(6’)-Ib7*, and *AAC(6’)-Ib9*) are here mostly found in integrons, agreeing with previous description of this class of ARGs^4^. The sulfonamide resistance *sul* ARO (3000410), which is canonically associated with Class 1 integrons, are actually only found in association with integrons in 10.49% of the total identified CRL occurrences (n = 1,173), while it is located outside of integrons in 89.51% of its CRLs.

**Supplementary Table 4.**
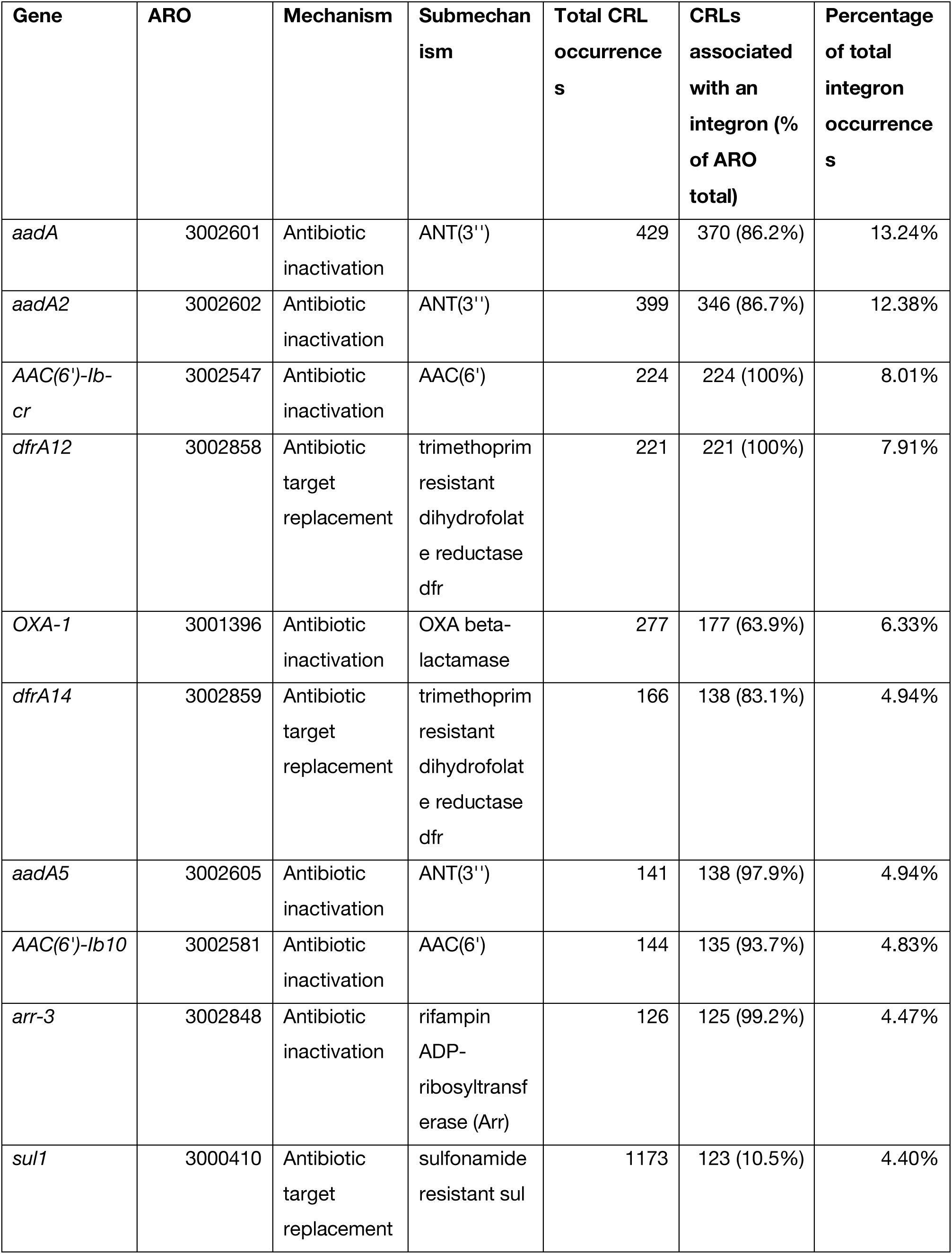
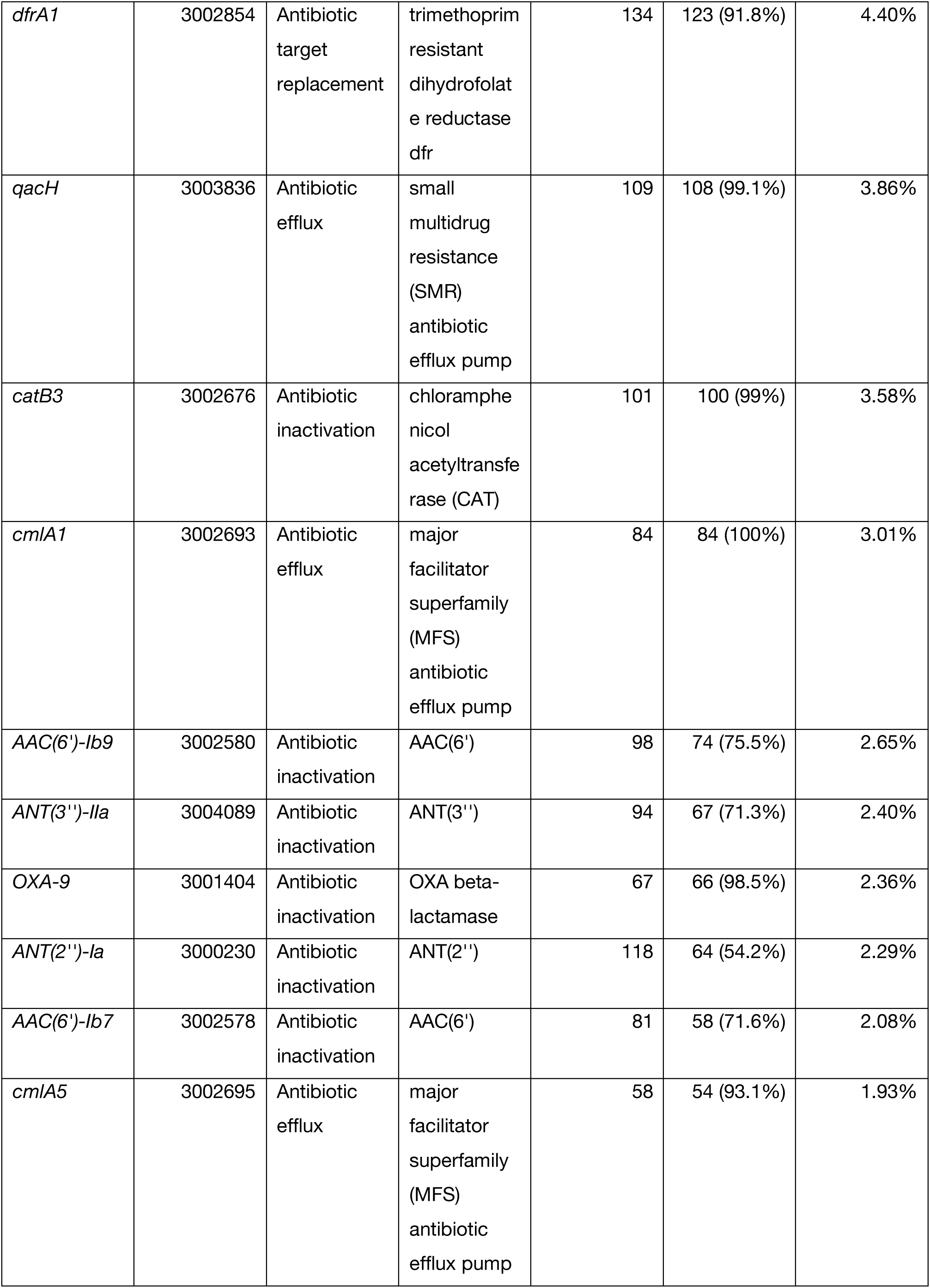
The top 20 AROs associated with integrons.

| Gene | ARO | Mechanism | Submechanism | Total CRL occurrences | CRLs associated with an integron (% of ARO total) | Percentage of total integron occurrences |
| --- | --- | --- | --- | --- | --- | --- |
| *aadA* | 3002601 | Antibiotic inactivation | ANT(3'') | 429 | 370 (86.2%) | 13.24% |
| *aadA2* | 3002602 | Antibiotic inactivation | ANT(3'') | 399 | 346 (86.7%) | 12.38% |
| *AAC(6')-Ib-cr* | 3002547 | Antibiotic inactivation | AAC(6') | 224 | 224 (100%) | 8.01% |
| *dfrA12* | 3002858 | Antibiotic target replacement | trimethoprim resistant dihydrofolate reductase dfr | 221 | 221 (100%) | 7.91% |
| *OXA-1* | 3001396 | Antibiotic inactivation | OXA beta-lactamase | 277 | 177 (63.9%) | 6.33% |
| *dfrA14* | 3002859 | Antibiotic target replacement | trimethoprim resistant dihydrofolate reductase dfr | 166 | 138 (83.1%) | 4.94% |
| *aadA5* | 3002605 | Antibiotic inactivation | ANT(3'') | 141 | 138 (97.9%) | 4.94% |
| *AAC(6')-Ib10* | 3002581 | Antibiotic inactivation | AAC(6') | 144 | 135 (93.7%) | 4.83% |
| *arr-3* | 3002848 | Antibiotic inactivation | rifampin ADP-ribosyltransferase (Arr) | 126 | 125 (99.2%) | 4.47% |
| *sul1* | 3000410 | Antibiotic target replacement | sulfonamide resistant sul | 1173 | 123 (10.5%) | 4.40% |
| *dfrA1* | 3002854 | Antibiotic target replacement | trimethoprim resistant dihydrofolate reductase dfr | 134 | 123 (91.8%) | 4.40% |
| *qacH* | 3003836 | Antibiotic efflux | small multidrug resistance (SMR) antibiotic efflux pump | 109 | 108 (99.1%) | 3.86% |
| *catB3* | 3002676 | Antibiotic inactivation | chloramphenicol acetyltransferase (CAT) | 101 | 100 (99%) | 3.58% |
| *cmlA1* | 3002693 | Antibiotic efflux | major facilitator superfamily (MFS) antibiotic efflux pump | 84 | 84 (100%) | 3.01% |
| *AAC(6')-Ib9* | 3002580 | Antibiotic inactivation | AAC(6') | 98 | 74 (75.5%) | 2.65% |
| *ANT(3'')-IIa* | 3004089 | Antibiotic inactivation | ANT(3'') | 94 | 67 (71.3%) | 2.40% |
| *OXA-9* | 3001404 | Antibiotic inactivation | OXA beta-lactamase | 67 | 66 (98.5%) | 2.36% |
| *ANT(2'')-Ia* | 3000230 | Antibiotic inactivation | ANT(2'') | 118 | 64 (54.2%) | 2.29% |
| *AAC(6')-Ib7* | 3002578 | Antibiotic inactivation | AAC(6') | 81 | 58 (71.6%) | 2.08% |
| *cmlA5* | 3002695 | Antibiotic efflux | major facilitator superfamily (MFS) antibiotic efflux pump | 58 | 54 (93.1%) | 1.93% |

### Supplementary Note 9: Pearson correlation coefficient analyses for major mechanisms

Per major mechanism, Pearson correlation coefficients were also calculated pairwise between each of the four MOB parameters (Supplementary Figs. 16-20). For all mechanisms, IS and Replicon ratios correlate with similar correlation coefficients. ARG-IS association likewise correlates with ARG- integron association, except for the ATA mechanism. For mechanisms AI and AE, all correlations are positive and significant. For ATR, the Simpson index correlates with neither IS nor integron ratio. For ATA and ATP, four and two correlations were not significant, respectively. Numerically, AE CRLs are most often found on chromosomes with no association with IS elements or integrons (Supplementary Fig. 21). Contrary to this, AI and ATR CRLs are very often found on plasmids in association with both IS elements and integrons. Overall, there is also an evident association between integrons and IS elements and integrons without IS elements is a rare combination.

**Supplementary Fig. 16.**
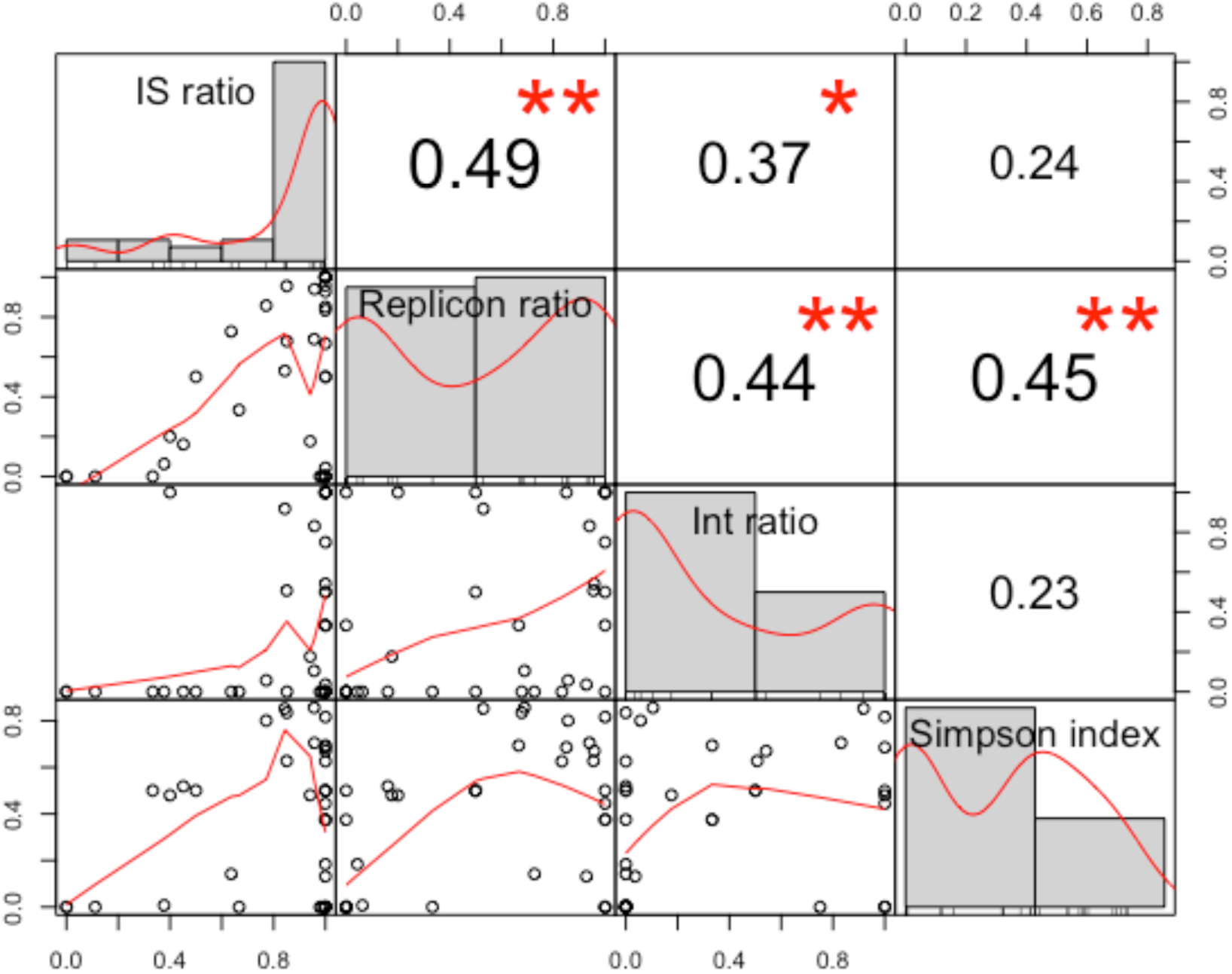
Pearson correlation coefficients for Antibiotic target replacement (ATR)

**Supplementary Fig. 17.**
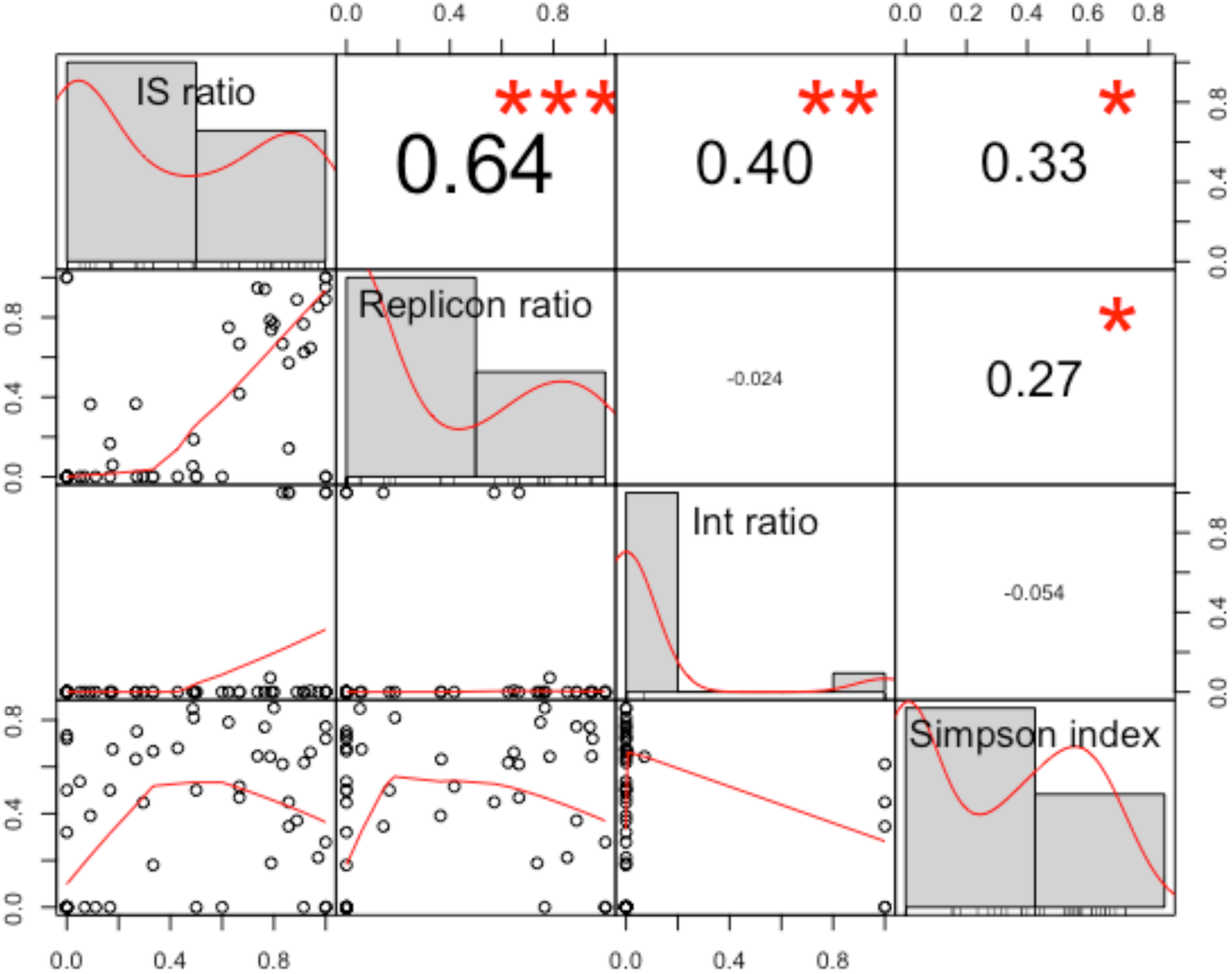
Pearson correlation coefficients for Antibiotic target protection (ATP)

**Supplementary Fig. 18.**
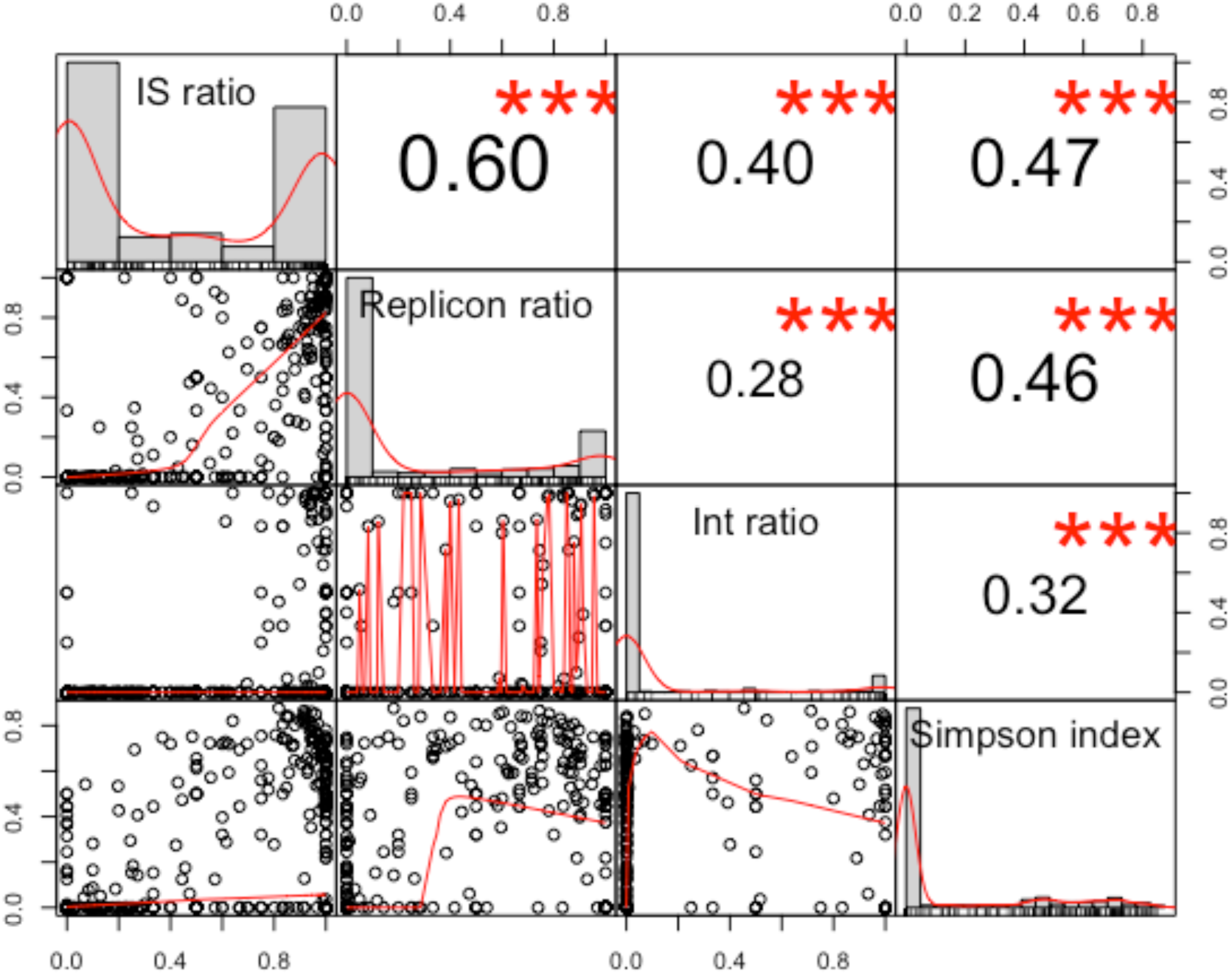
Pearson correlation coefficients for Antibiotic inactivation (AI)

**Supplementary Fig. 19.**
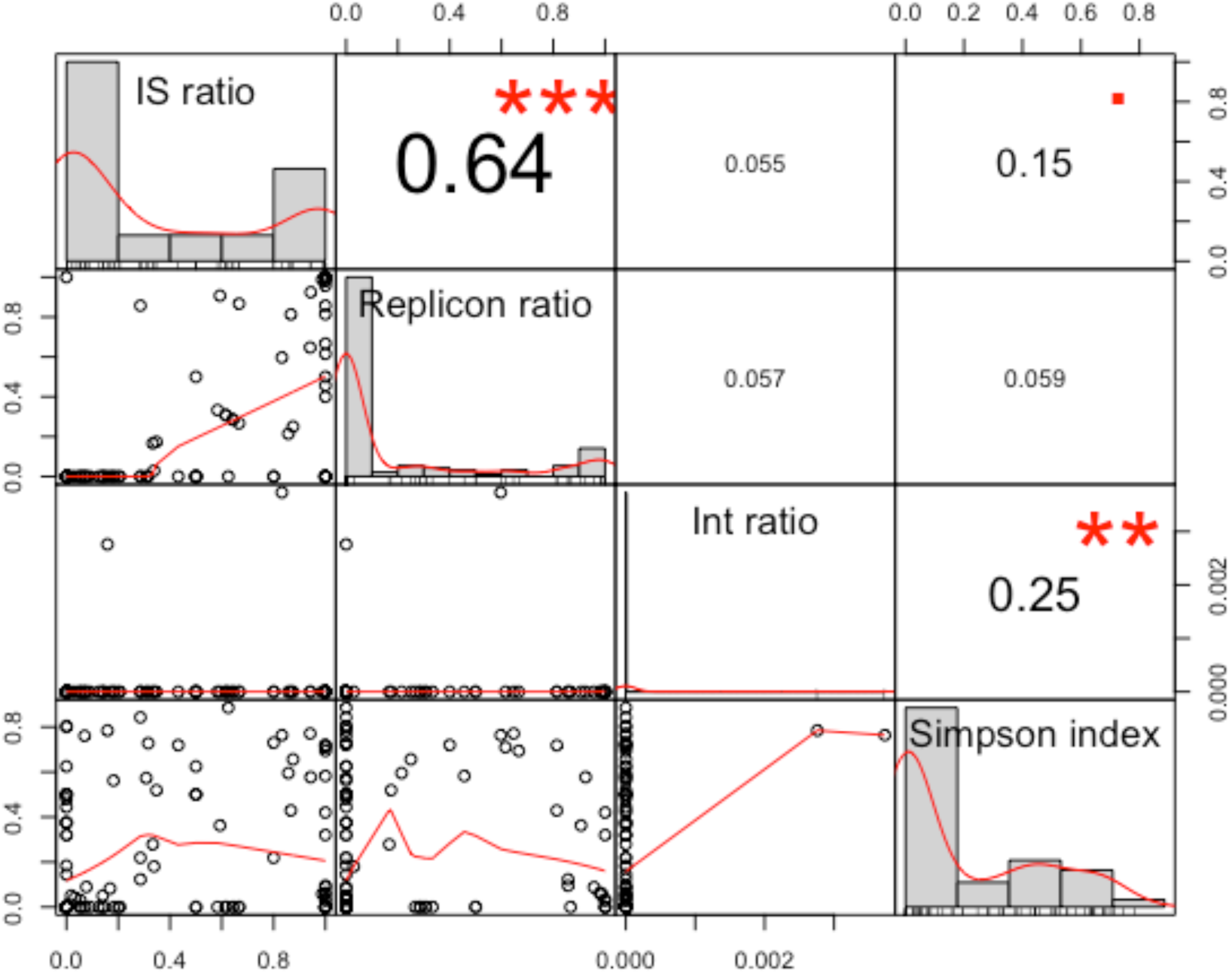
Pearson correlation coefficients for Antibiotic target alteration (ATA)

**Supplementary Fig. 20.**
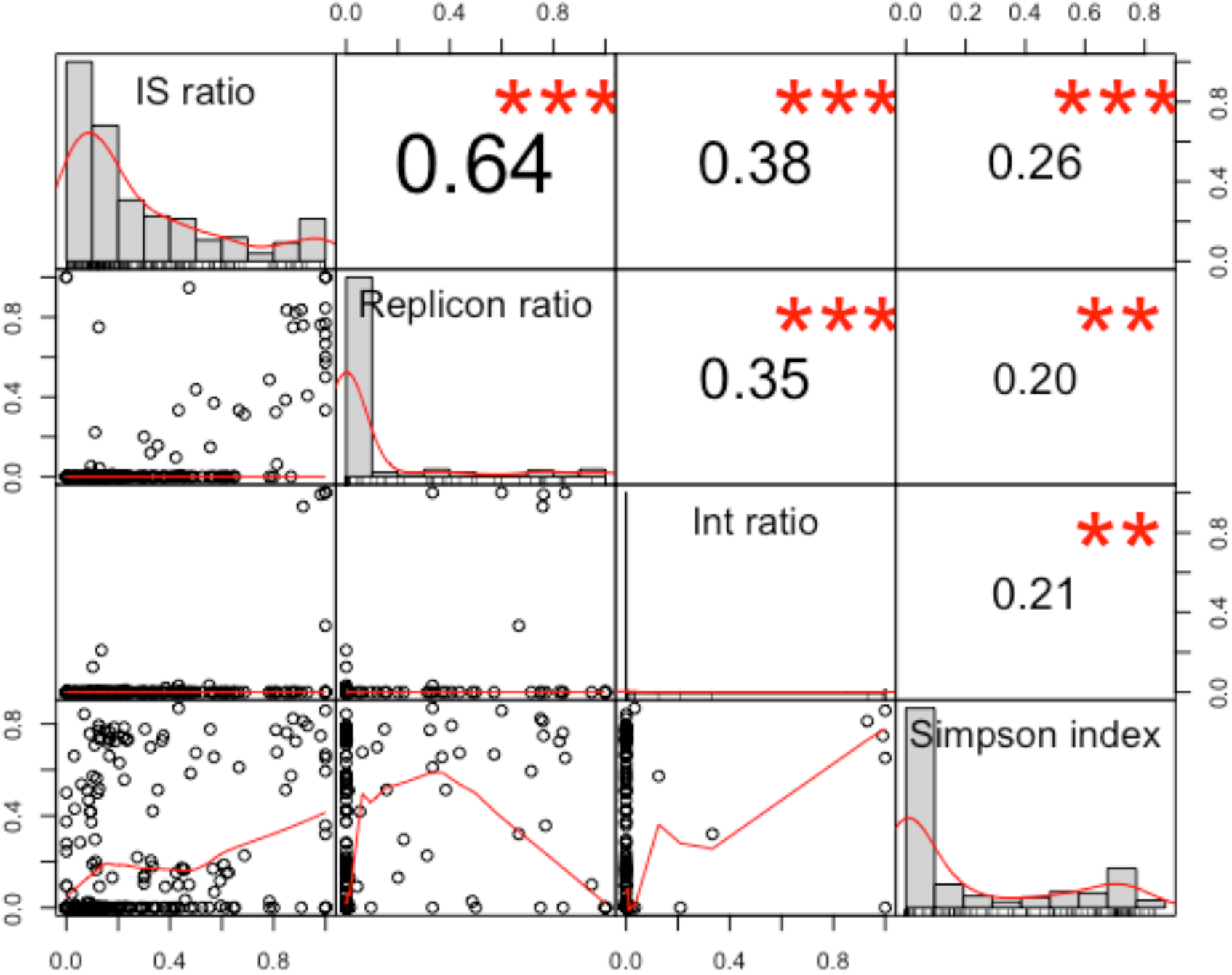
Pearson correlation coefficients for Antibiotic efflux (AE)

**Supplementary fig. 21:**
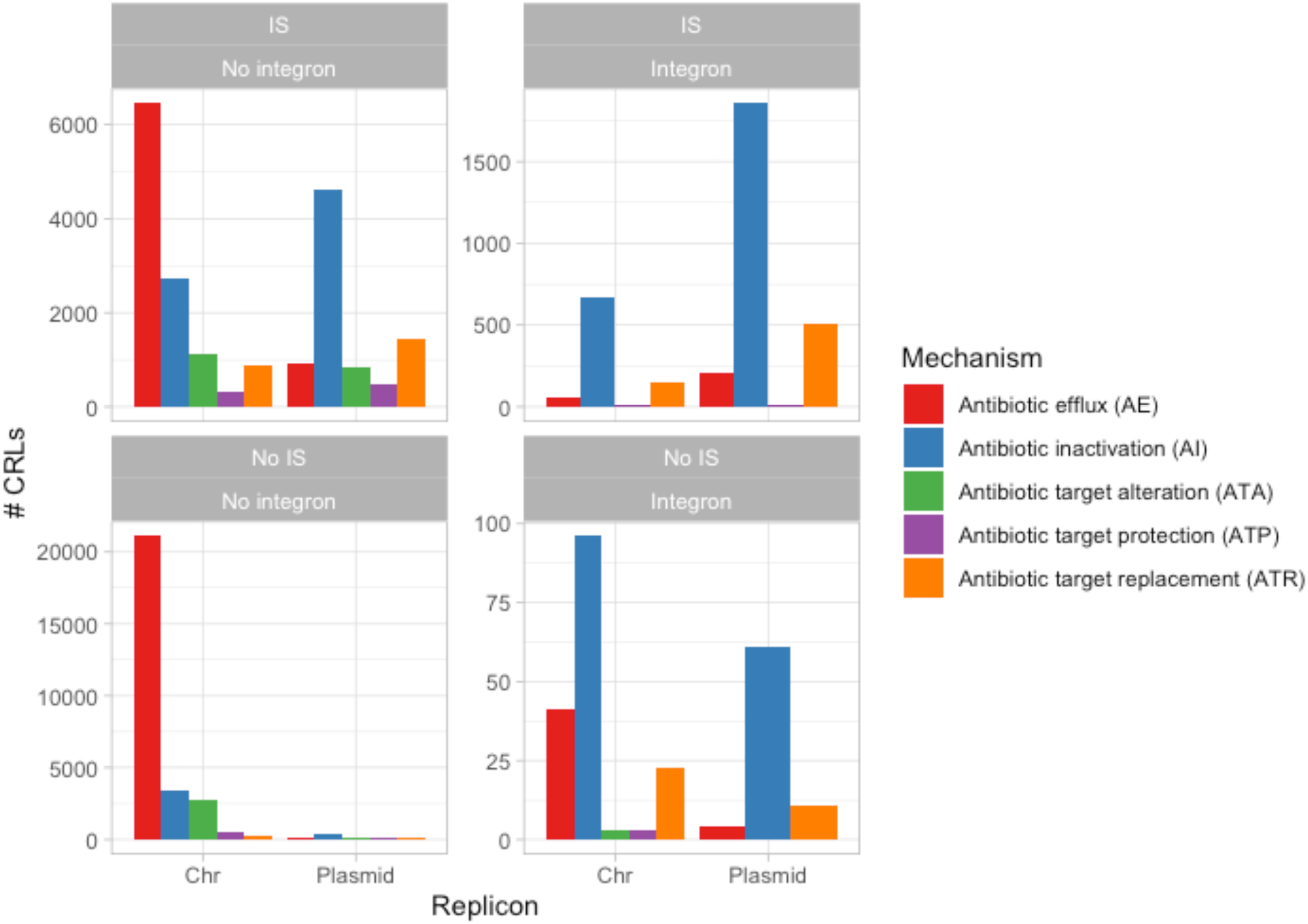
Counts of CRLs separated by replicon type, and IS- and integron association. Bars are colored by resistance mechanism. Y-axis scales are different between each subplot.

### Supplementary Note 10: Some AROs are highly divergent in mobilization

The mean IS and Replicon ratios per ARO are calculated across all genera they are found in. However, upon closer scrutiny of AROs per genus it becomes obvious that some AROs have a high spread from their IS and Replicon ratio means. For Supplementary Fig. 22, the genus-specific ratios were calculated per ARO and their difference from the mean global ARO ratio was calculated per genus. For each ARO, the summed positive and negative differences (for each genus with a given ARO) are shown below. A positive difference from the ARO mean indicates that there are genera in which ARGs of the given ARO are more mobilized than the ARO mean. Vice versa, a negative summed difference from the mean shows that ARGs of a given ARO are less mobilized in some genera than the ARO mean.

**Supplementary Fig. 22:**
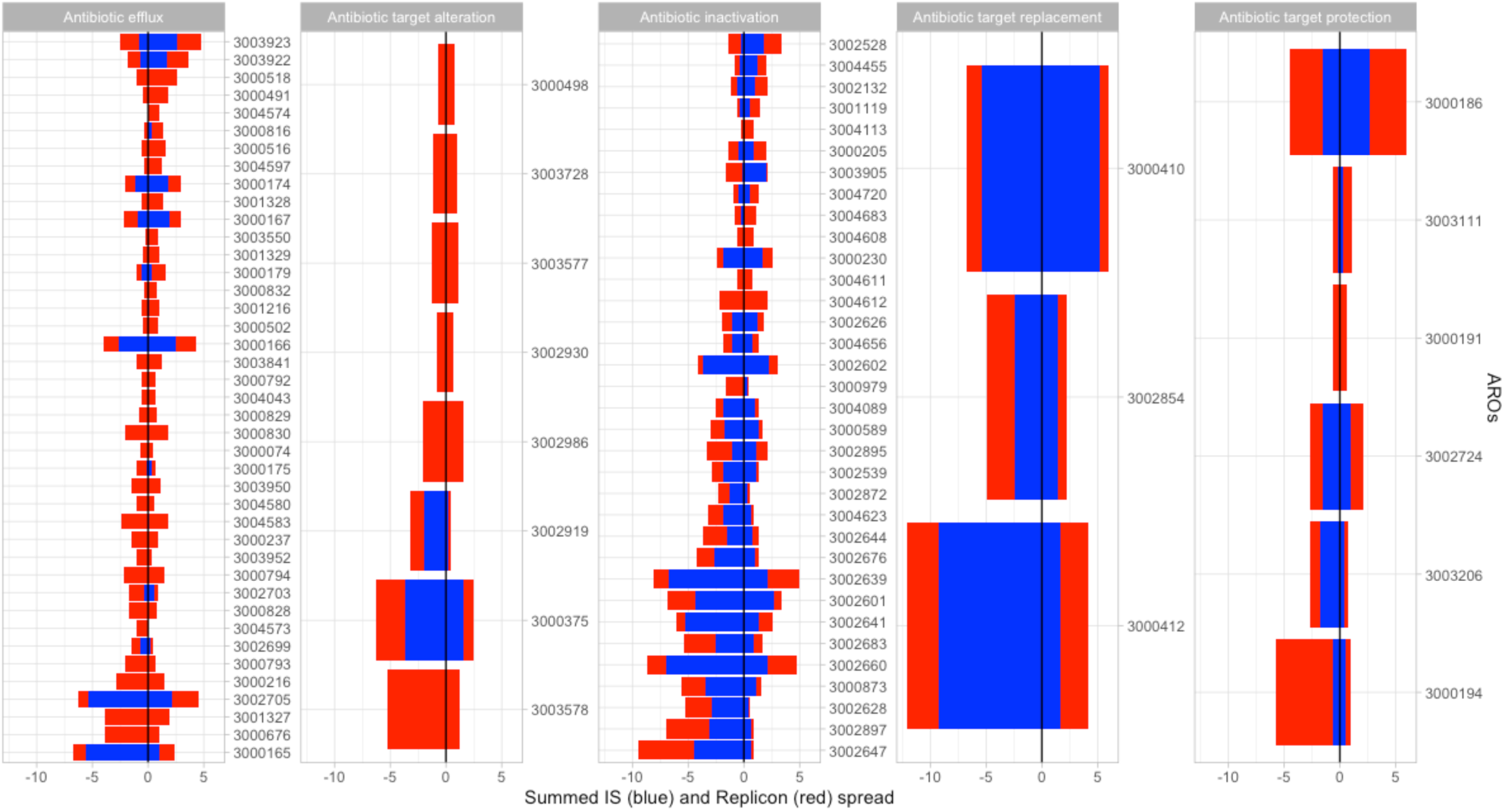
Summed spread from mean IS and Replicon ratios from the ARO means. Differences from the mean per ARO were summed by the genera in which the AROs were identified. A positive summed difference from ARO mean indicates that some genera have more mobilized ARGs of a given ARO than the ARO mean. Mobilization by both IS elements (left) and plasmids (right) are shown. Before calculating the genus-specific IS and Replicon ratio per ARO, genus:ARO combinations with only 1 occurrence in the dataset were filtered, since they are a source of noise in this context. Only AROs with a summed negative plus positive spread from mean of at least 1 are shown.

From the same dataset, genera are plotted with their summed spread from the mean of all AROs found within the respective genera (Supplementary Fig. 23). Especially AI AROs in *Shigella* are highly mobilized by IS elements compared to the ARO means. Interestingly, the AI AROs are not very mobilized by plasmids in *Shigella*, indicating that AI ARGs often associated with IS elements in *Shigella* but mostly on chromosomes. Also worth noting, *Proteus*, *Pseudomonas*, *Morganella*, *Acinetobacter*, and a few other genera have large negative summed Replicon spreads from mean, indicating that chromosomes in these may act as reservoirs for yet unmobilized ARGs. On the other hand, these putative and potential ARGs are not decontextualized and will likely occur as false-positive resistance genes in studies applying (q)PCR and metagenomic sequencing in environmental samples with these bacteria.

**Supplementary Fig. 23:**
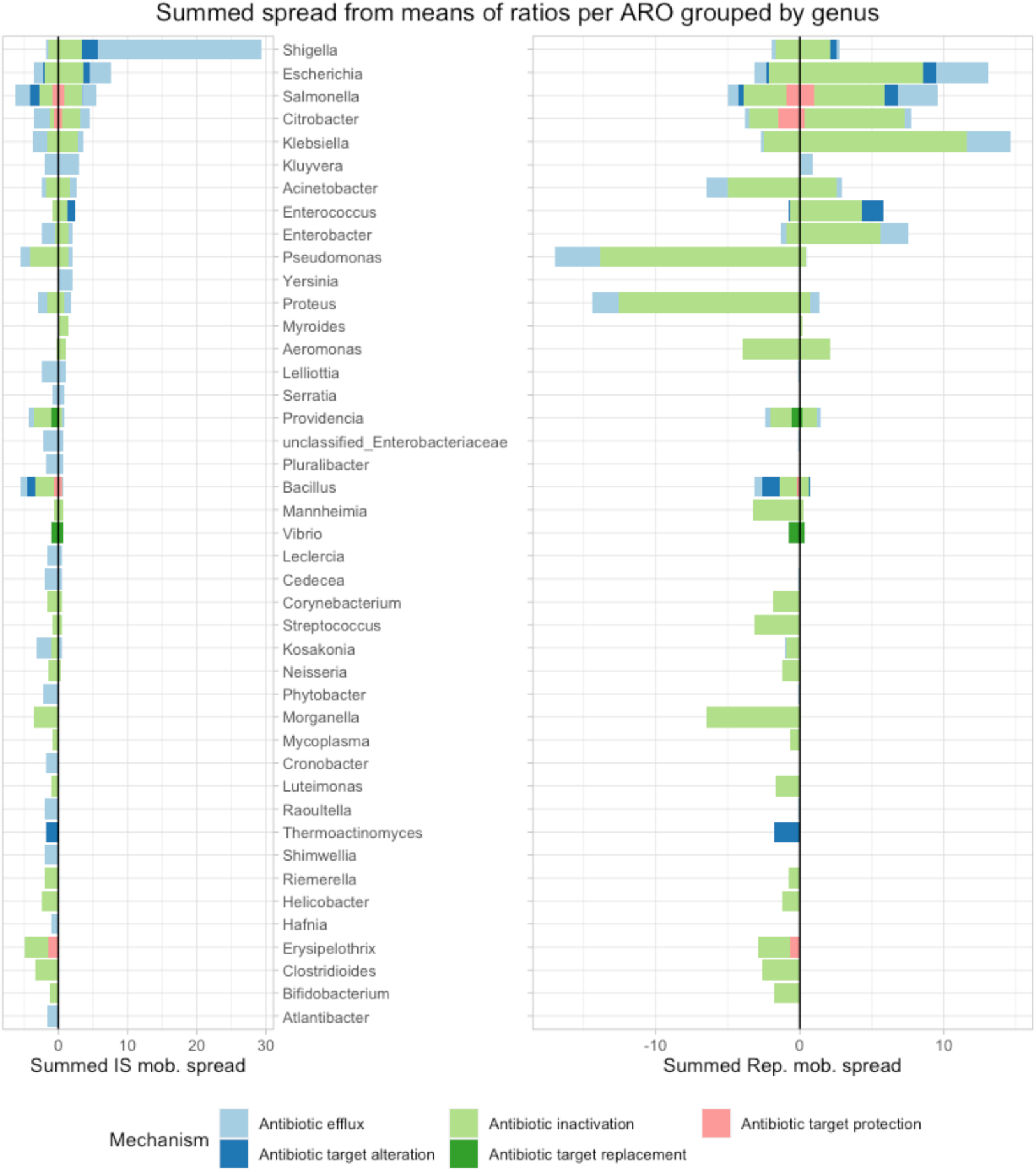
Summed spread from mean IS and Replicon ratios from the ARO means grouped by genera. The same dataset as for figure Supplementary Fig. 22 is used here, but the summed spreads are grouped by genera. A large positive summed spread indicates that a given genus has AROs that are more mobilized, by either IS elements or plasmids, than the ARO mean.

In Supplementary Table 5, an example of an ARO that is highly differential in IS and plasmid mobilization, depending on the genus it is found in. The OqxAB efflux pump is encoded by the neighboring genes *oqxA* and *oqxB* that have the same mobilization characteristics. Only results for *oqxA* are shown in this example, since *oqxB* has similar results (not shown). In *Klebsiella* genomes, *oqxA* is found in association with IS elements in 15.5% of cases and it is only on plasmids in 0.2%. As such, it should be considered a housekeeping genes, since it does not usually confer resistance unless highly overexpressed^5–7^.

**Supplementary Table 5.**
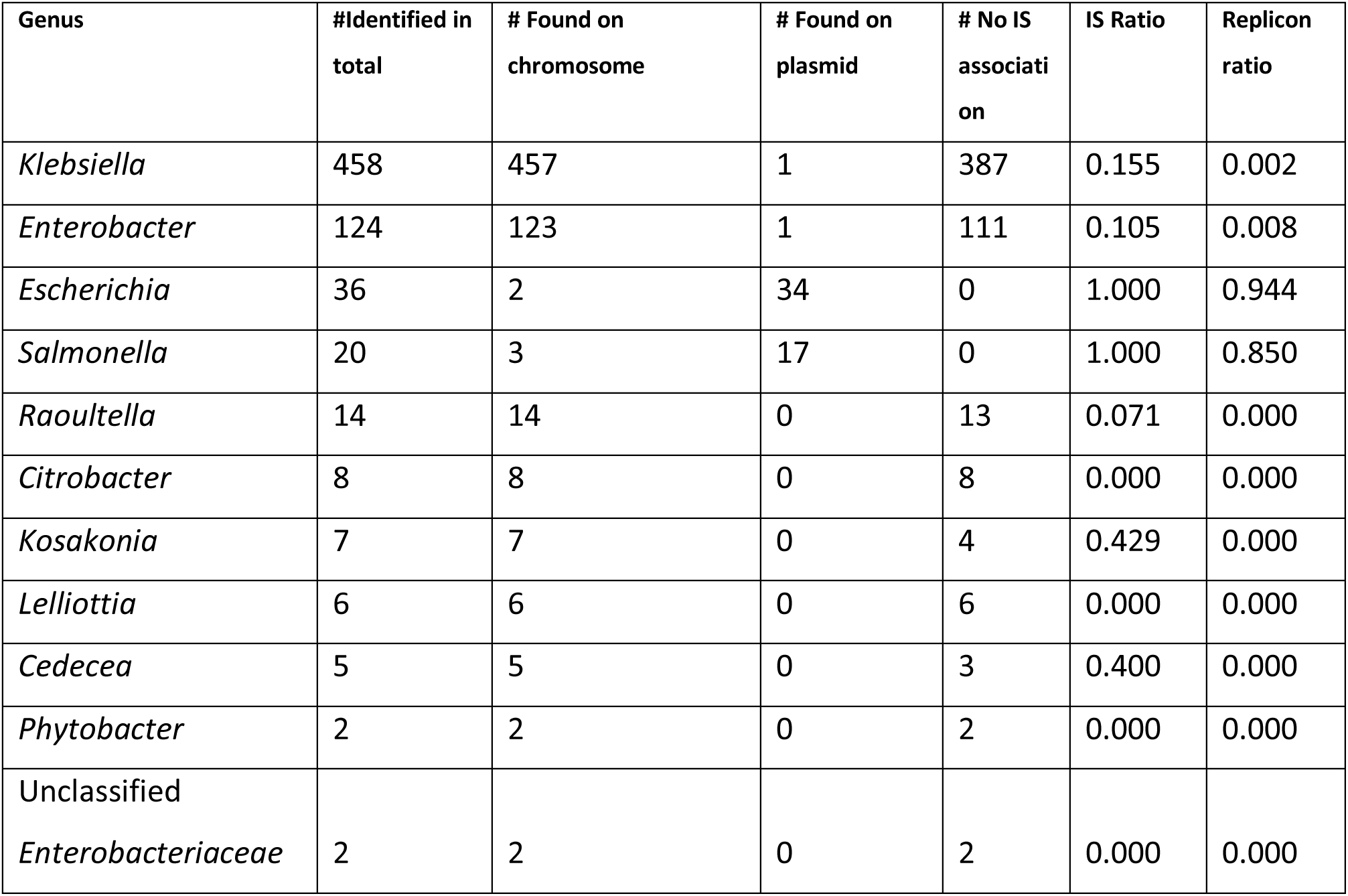
Information about the *oqxA* ARO.

**Supplementary Fig. 24:**
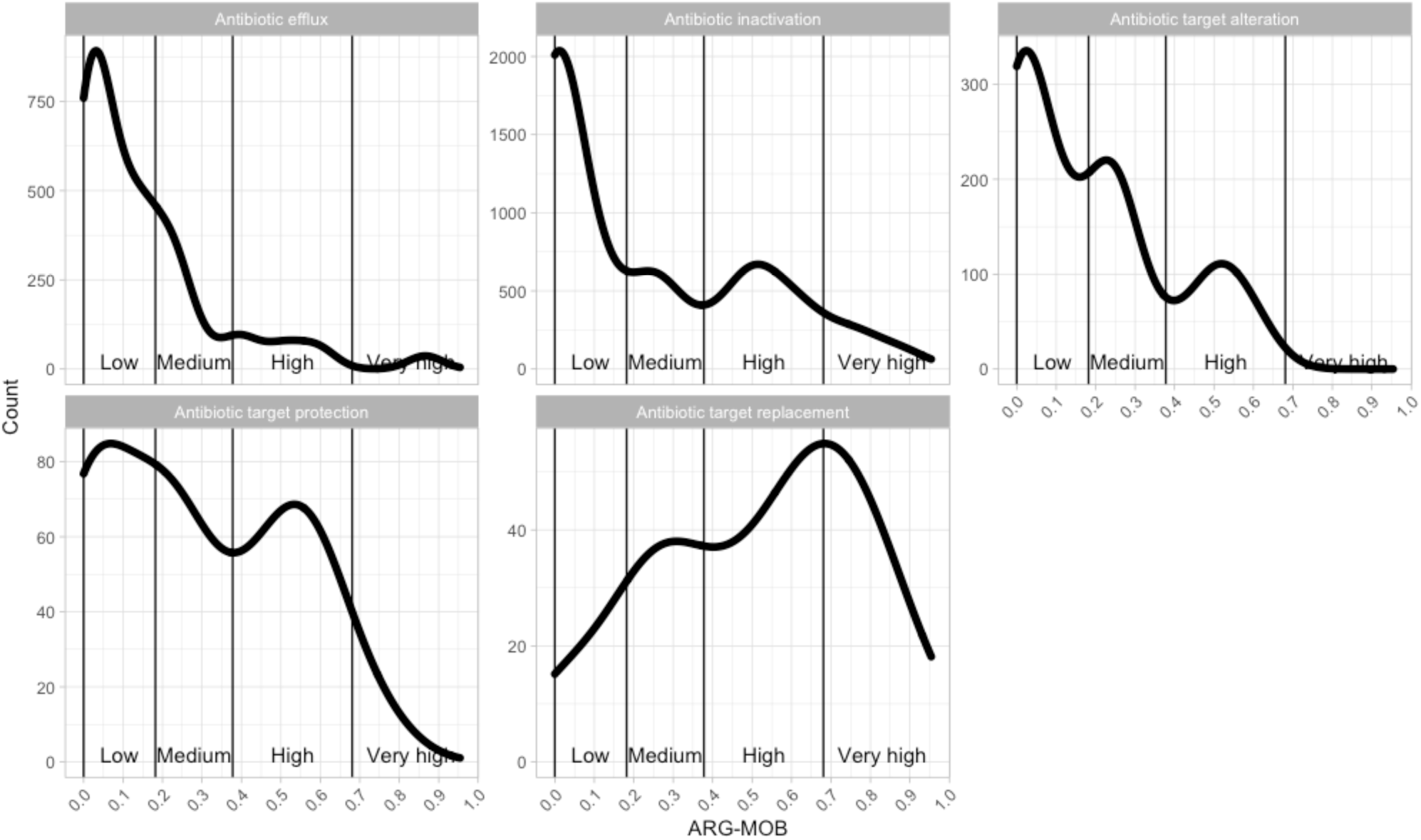
Count density of ARG-MOB per mechanism. Y-axes are not on the same scale between subplots. The global ARG-MOB category definitions are shown with vertical lines.

### Supplementary Note 11: Examples of *Low* and *High* ARG-MOB AROs

#### *Low* ARG-MOB examples

**ARG-MOB=0; 3003785 (ATA) MurA transferase:** Shown to confer resistance when cloned onto pBAD18 under overexpression in *E. coli* from *Chlamydia*^8^

**ARG-MOB=0; 3000421 (AE) NorB major facilitator superfamily (MFS) antibiotic efflux pump:** Requires overexpression for resistance to quinolones, tetracycline and others. Was cloned from *Staphylococci* into *E. coli* for overexpression to give resistance^9^.

**ARG-MOB=0; 3000462 (AI) MgtA mgt macrolide glycotransferase resistance:** Was shown to confer resistance to macrolides in *Streptomyces lividans* under overexpression from multicopy plasmid pLST21 with constitutive expression^10^.

**ARG-MOB=0; 3004143 (AE) resistance-nodulation-cell division (RND) antibiotic efflux pump:** The RND efflux pump AxyXY-OprZ is repressed by upstream transcriptional repressor axyZ. Gene deletion mutation of axyZ leads to increased axyZ transcription and increased MICs of multiple antibiotics (fluoroquinolones, cefepime, tetracyclines) in *Achromobacter*^11^. The AxyXY are orthologs of MexX and MexY in *Pseudomonas aeruginosa* where mutations in the mexZ transcriptional regulator leads overexpression of the MexXY efflux pump and increased resistance^12^. These pumps are likely naturally occurring pumps that require significant overexpression to yield problematic resistance.

**ARG-MOB=0; 3004775 (AI) CME beta-lactamase:** CME-1 was cloned from a *Flavobacterium* and overexpressed from a cloning vector in *E. coli* to provide resistance to cefuroxime^13^. The strain was not a clinical isolate.

**ARG-MOB=0; 3003035 (ATP) MfpA quinolone resistance protein (qnr):** MfpA from *Mycobacterium tuberculosis* strain H37Rv protects DNA gyrase from fluoroquinolone inhibition. It was found to confer low-level resistance when overexpressed from a multicopy plasmid^14^, but strain H37Rv is not inherently resistant to fluoroquinolones^15^.

**ARG-MOB=0; 3002646 (AI) APH(3’) aminoglycoside phosphotransferase:** Aph(3’)-IIc was cloned and overexpressed from the chromosome of *Stenotrophomonas maltophilia* in *E. coli* to confer increased MICs to amikacin, butirosin, kanamycin, lividomycin, neomycin, paromomycin, and tobramycin. In this study, Aph(3’)-IIc was knocked out on the chromosome of the WT strain and MICs to butirosin, kanamycin, neomycin and paromomycin were decreased, showing the this gene indeed confers intrinsic resistance those antibiotics. While expression of Aph(3’)-IIc was evaluated with qPCR in the WT strain, the expression levels were not reported and it is thus not known if resistance from this gene is due to mutation in an expression regulator gene in the clinical *Stenotrophomonas* isolate^16^.

#### *High* ARG-MOB examples

**ARG-MOB=0.95; 3003013 (ATR) *dfrA15* (trimethoprim resistant dihydrofolate reductase dfr):** A Class 1 integron with *dfrA15* is widespread in *Vibrio cholera* isolates in Africa, causing resistance to trimethoprim. It was furthermore found on a conjugative plasmid^17^. In this study, it is the ARO with the highest risk, due to the fact that it was only found to be associated with IS elements, integrons and plasmids (all ratios = 1). It has a Simpson index of 0.82 and the 7 CRLs are dispersed across 6 genera (*Vibrio*, *Salmonella*, *Enterobacter*, *Leclercia*, *Klebsiella*, and *Escherichia*).

**ARG-MOB=0.91; 3002271 (AI) VIM-1 (VIM beta-lactamase):** VIM-1 was originally isolated from a multiresistant *E. coli* from a patient in Greece. It was inserted in a class 1 integron and found on a conjugative plasmid^18^. It has since been seen in multiple *Enterobacteriaceae*, typically in association with integrons and plasmids, and is globally spread. Here, it scores the fourth-highest ARG-MOB= of 0.91 with IS, replicon, and integron ratios of 0.95, 0.95, and 1, respectively. The 21 CRLs are found in 6 distinct genera (*Pseudomonas*, *Salmonella*, *Escherichia*, *Klebsiella*, *Citrobacter*, and *Enterobacter*).

**ARG-MOB=0.89; 3004635 (AI) AAC(6’):** During an *Enterobacter* outbreak in Venezuela, the multi-resistance encoding conjugative plasmid pBWH301 was isolated. Amongst other ARGs, *aacA7* was found to encode AAC(6’)-I aminoglycoside acetyltransferase in an integron^19^. In RefSeq complete genomes, there are 12 CRLs which are all on plasmids and associated with IS elements. Furthermore, it is inserted in integrons in 92% of cases. It is dispersed across 3 genera for a Simpson index of 0.63. Several other AROs for AAC(6’) subtypes are among the highest ARG-MOB scoring AROs, resulting in one of the highest mean ARG-MOB scores for any Antibiotic Inactivation submechanism ().

**ARG-MOB=0.86; 3002847 (AI) rifampin ADP-ribosyltransferase (Arr):** From a multi-resistant clinical *P*. *aeruginosa* isolate, a class I integron was cloned into an expression vector and transformed into *E. coli* to screen for rifampin resistance^20^. Although an expression vector was used to identify rifampin resistance genes under heterologous expression, it was described that the DNA insert of the clone carrying rifampin resistance gene *arr-2* also carried a class I integron with *arr-2* inserted as a gene cassette. This corroborates our finding that *arr-2* is a high risk putative ARG where it was associated with IS elements in 94% of CRLs, found on plasmids on 87%, and found as integron gene cassettes in 96% of CRLs (n=54). Furthermore, it was found in 8 distinct genera, for a Simpson index of 0.64 (*Acinetobacter*, *Citrobacter*, *Escherichia*, *Klebsiella*, *Proteus*, *Pseudomonas*, *Salmonella*, and *Shewanella*).

